# Retinal microvascular and neuronal pathologies probed *in vivo* by adaptive optical two-photon fluorescence microscopy

**DOI:** 10.1101/2022.11.23.517628

**Authors:** Qinrong Zhang, Yuhan Yang, Kevin J. Cao, Wei Chen, Santosh Paidi, Chun-Hong Xia, Richard H. Kramer, Xiaohua Gong, Na Ji

## Abstract

The retina, behind the transparent optics of the eye, is the only neural tissue whose physiology and pathology can be non-invasively probed by optical microscopy. The aberrations intrinsic to the mouse eye, however, prevent high-resolution investigation of retinal structure and function *in vivo*. Optimizing the design of a two-photon fluorescence microscope (2PFM) and sample preparation procedure, we found that adaptive optics (AO), by measuring and correcting ocular aberrations, is essential for resolving synapses and achieving three-dimensional cellular resolution in the mouse retina *in vivo*. Applying AO-2PFM to longitudinal retinal imaging in transgenic models of retinal pathology, we characterized microvascular lesions and observed microglial migration in a proliferative vascular retinopathy model, and found Lidocaine to effectively suppress retinal ganglion cell hyperactivity in a retinal degeneration model. Tracking structural and functional changes at high resolution longitudinally, AO-2PFM enables microscopic investigations of retinal pathology and pharmacology for disease diagnosis and treatment *in vivo*.

## Introduction

Retina is a layered tissue in the back of the eye that transduces light into electrochemical signals to be further processed by the brain for visual perception and cognition^1^. As one of the most energy-demanding tissues, the retina is metabolically sustained by an intricate vasculature with several laminar plexuses^2^. Vascular and neuronal abnormalities in the retina are associated with both ocular^3^ and systemic diseases^4–6^, underscoring the importance of studying retinal pathology and pharmacology.

With well-developed genetics and similar physiology to the human retina, mouse models have been widely utilized for mechanistical studies of retinal diseases. Behind highly transparent mouse eye optics (i.e., cornea and crystalline lens), the retina is uniquely accessible to light and the only part of the nervous system that can be probed non-invasively by optical imaging. Recent advances in mouse genetics have enabled fluorescence microscopy investigations of vasculature^7^ as well as neurons and glial cells^8–10^ of the mouse retina. Among fluorescence microscopy techniques, two-photon fluorescence microscopy (2PFM)^11^ utilizing near-infrared (NIR) excitation is particularly suited for retinal imaging. Its intrinsic optical sectioning capability permits depth-resolved three-dimensional (3D) imaging throughout the retina. With the retinal photoreceptors minimally responsive to NIR light, 2PFM is also an ideal tool for functional studies of retina^12,13^. However, as a far-from-perfect imaging system, the optics of the mouse eye introduce severe aberrations to the NIR excitation light, preventing high-resolution visualization of subcellular features *in vivo*. As a result, the vast majority of microscopy studies have been carried out *ex vivo* on dissected retinas, preventing longitudinal investigations of retinal pathology under physiological conditions.

Adaptive optics (AO) is a collection of technologies that actively measure and correct for optical aberrations^14^, and has been applied to optical microscopy for high-resolution imaging of neural tissues^15,16^. It has also been combined with ophthalmological imaging modalities to restore diffraction-limited imaging performance for the human retina^17,18^. Because of the severe aberrations of the mouse eye, AO has also been applied to *in vivo* imaging of the mouse retina^19–26^. However, there are disagreements in the reported spatial resolutions^19–26^, characteristics and magnitude of aberration^19–21,24–26^, and the effectiveness of AO^20,21,24–26^. For example, whereas previous papers reported cellular resolution without AO, a recent AO-2PFM study^26^ reported extremely large aberrations in the mouse eye and found AO to be required in order to resolve microvasculature and cell bodies in 2D *in vivo*. These discrepancies have led to uncertainty over the imaging performance achievable with conventional 2PFM and the necessity of AO for microvascular and cellular investigations of retinal physiology. Together with a lack of detailed imaging protocols, they have prevented the routine application of AO-2PFM to disease diagnosis and therapeutic intervention in the retina of mouse models of ocular, cerebral, and systemic diseases.

The aims of this work are to provide a resource for *in vivo* retinal imaging using 2PFM, by optimizing the design of a 2PFM for *in vivo* imaging of the mouse retina, characterizing mouse ocular aberration, developing a guideline for adaptive optical 2PFM (AO-2PFM) imaging, and demonstrating its applications to retinal pathology and pharmacology. Using a carefully engineered 2PFM and following an optimized sample preparation procedure, we were able to achieve two-dimensional (2D) cellular resolution imaging performance in the mouse retina without AO. For synaptic, subcellular, and three-dimensional (3D) cellular resolution imaging of the mouse retina, AO was essential in improving image brightness, contrast, and resolution. Testing the performance of AO-2PFM in various transgenic mouse lines, we proposed strategies to maximize its impact on image quality improvement. We extended the application of AO-2PFM to mouse retinal pathology and pharmacology by imaging the retinas of two transgenic models with proliferative vascular retinopathy and retinal degeneration, respectively. In our model of proliferative vascular retinopathy, AO enabled us to, for the first time, characterize retinal vascular lesions with sub-capillary details over multiple days and track microglial migration *in vivo*. In our model of retinal degeneration, AO allowed high-fidelity interrogation of pharmacologically modified hyperactivity of retinal ganglion cells (RGCs), indicating AO-2PFM as a promising tool evaluating retinal pharmacology *in vivo*. Together, by systematically optimizing and applying AO-2PFM to *in vivo* mouse retinal imaging, our work represents an important advancement in enabling high-resolution longitudinal studies of retinal pathology and pharmacology for disease diagnosis and treatment.

## Results

### Optimized AO-2PFM for *in vivo* mouse retinal imaging

A home-built 2-photon fluorescence microscope equipped with a segmented deformable mirror (DM) and a Shack-Hartmann (SH) sensor^27^ was modified for *in vivo* mouse retinal imaging by replacing the objective lens with an add-on eye imaging module^26,28^ (**Fig. 1A, Materials and Methods**). The module consisted of an electrically tunable lens (ETL) whose adaptive surface was conjugated to the DM, a turning mirror, and two lens groups (L7 and L8) that relayed the adaptive surface of the ETL to the pupil of the mouse eye. With this design, the optics of the mouse eye focused 920 nm light onto the retina to excite fluorescent markers and collected the emitted fluorescence for detection. The ETL allowed us to adjust the focal plane in the mouse eye without translating the mouse^29^ or optics^30^ in the imaging system. For all experiments, system aberrations in the 2-photon illumination path were measured with a modal AO method and corrected before image acquisition (**Materials and Methods**; “No AO” images: system aberration correction only).

**Figure 1.**
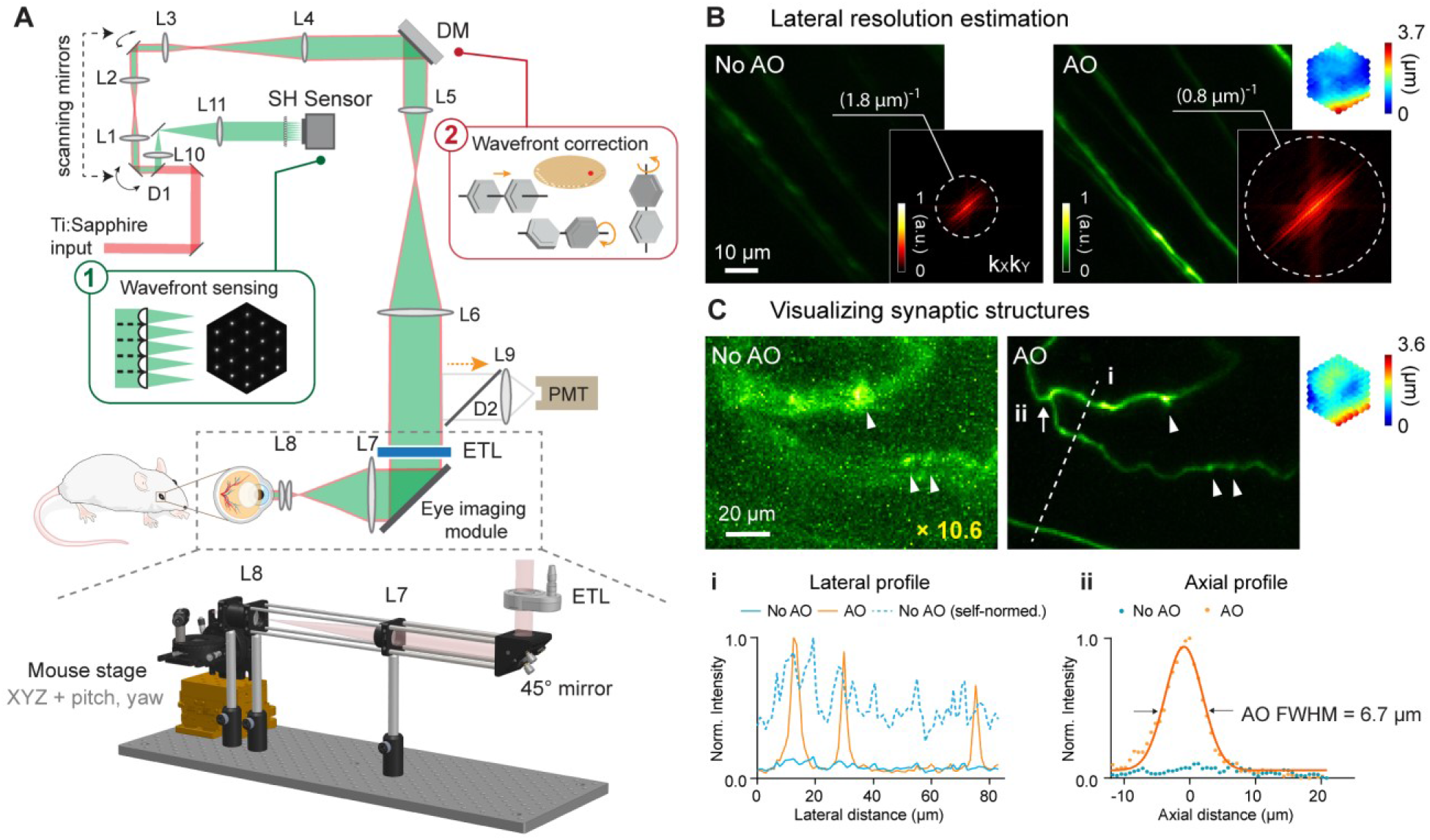
AO-2PFM for diffraction-limited imaging of the mouse retina *in vivo*. **(A)** Schematics of AO-2PFM. Inset 1: direct wavefront measurement by a Shack-Hartmann (SH) sensor composed of a lenslet array and a camera. Inset 2: wavefront correction with a deformable mirror composed of 163 segments with piston, tip, and tilt controls. Grey dashed box: eye imaging module. Bottom: 3D assembly of eye imaging module. L, lens; D, dichroic mirror; DM, deformable mirror; PMT, photomultiplier tube; ETL, electrically tunable lens. **(B)** Maximum intensity projections (MIPs) of image stacks (72 × 72 × 25 µm3) of RGC axons measured without and with AO, respectively, normalized to AO image. Insets: kXkY spatial frequency representation of the images and corrective wavefront. **(C)** MIPs of image stacks (132 × 97 × 32 µm^3^) of fine RGC processes measured without and with AO, respectively, normalized to AO image. ‘No AO’ image brightness artificially increased by 10.6× for better visualization. White arrowheads: synaptic structures. Inset: corrective wavefront. Bottom: i: lateral signal profiles along white dashed line; ii: axial signal profiles of process ii (white arrow). Representative data from > 3 experiments (technical replicates).

To ensure optimal performance, we thoroughly characterized our AO-2PFM. We investigated how ETL current and mouse eye placement (with a longitudinal displacement of up to 4 mm in typical experiments) impacted imaging performance (**Fig. S1**). We found that aberrations introduced by the ETL at different control currents minimally affected image quality and that axial focal shift varied linearly with ETL current while field-of-view (FOV) size remained mostly constant. We also optimized sample preparation procedure. We discovered that a custom-designed 0-diopter contact lens (CL; design parameters in **Fig. S2A**) in combination with a single application of eye gel between the CL and the cornea reduced aberrations, prevented cataract formation, and improved wavefront sensing and imaging for hours (**Fig. S2**).

In order to achieve diffraction-limited imaging of the mouse retina *in vivo*, we measured and corrected ocular aberrations with a direct wavefront sensing method^31,32^, utilizing the SH sensor for wavefront measurement and the DM for wavefront correction (**Fig. 1A**). Briefly, a 3D-localized fluorescence ‘guide star’ was formed in the retina via 2-photon excitation and scanned over a user-defined 2D area with galvanometer scanning mirrors. The emitted fluorescence was collected and, after being descanned by the same pair of scanning mirrors, directed to the SH sensor. The now stationary fluorescence wavefront was segmented by a lenslet array and focused onto a camera, forming an SH image composed of an array of foci (**Fig. 1A**, inset 1). Local phase slopes of wavefront segments were calculated from the displacements of the foci from those taken without aberrations. Assuming spatially continuous aberrations, we computationally reconstructed the wavefront from the phase slopes^33^. We then applied a corrective wavefront, opposite to the measured aberrations, to the DM by controlling the tip, tilt, and piston of each segment (**Fig. 1A**, inset 2) so that mouse ocular aberrations could be canceled out, ensuring diffraction-limited focusing of the 2-photon excitation light on the mouse retina.

All *in vivo* imaging experiments were conducted in anesthetized mice with dilated pupil (**Materials and Methods**). In most experiments, an area of 19 × 19 µm^2^ of the retina was scanned for 3-10 seconds for wavefront sensing. To estimate the spatial resolution of our AO-2PFM for *in vivo* mouse retinal imaging, we imaged Thy1-GFP line M transgenic mice that had green fluorescent protein (GFP) expressed in a subset of RGCs^34^. The image taken without AO showed dim and distorted RGC axons; after aberration correction, we achieved an 8.6× increase in signal and proper visualization of the fine RGC axons (**Fig. 1B**). The spatial frequency space representations of the images indicated that AO enhanced the ability of the imaging system to acquire higher resolution information and led to a lateral resolution that was better than ~0.8 µm (**Fig. 1B**, insets). For some thin RGC processes (**Fig. 1C**), restoring diffraction-limited resolution led to an increase in signal (by 10.6×) and contrast (**Fig. 1C, i**), and, for the first time, enabled *in vivo* 2PFM visualization of synaptic structures in the mouse retina (**Fig. 1C**, white arrows). From the axial profile of a thin process (**Fig. 1C, ii**), we estimated the axial resolution after AO correction to be 6.7 µm. Both the lateral and axial resolution estimations were close to the theoretical diffraction-limited resolution for a fully-dilated mouse eye with 0.49 numerical aperture^35^.

### AO improves *in vivo* imaging of retinal vasculature

Retinal vasculature supports the physiological functions of the retina. Retinal vascular diseases can lead to vision loss. Abnormalities in retinal vasculature morphology and physiology serve as important biomarkers for various cerebral and systemic diseases^36–39^. Therefore, *in vivo* characterization of retinal vasculature, especially at the microvasculature level, is of great physiological and clinical importance. Utilizing either confocal microscopy^20,25^ or 2PFM^26,40,41^, previous publications have achieved *in vivo* visualization of retinal microvasculature through either full correction of the mouse eye aberrations^20,25,26^, partial correction of the anterior optics of the mouse eye^41^, or stringent selection of imaging lenses^40^. These prior demonstration-of-principle experiments suggest that in order to image retinal microvasculature *in vivo*, mouse eye aberrations need to be corrected, either fully or partially. With our optimized imaging system, we aimed to determine whether aberration correction was indeed essential for visualizing microvasculature. Furthermore, we proceeded to systematically characterize the spatial dependence of mouse eye aberrations and how large a FOV can benefit from a single AO correction.

To verify the necessity of AO in resolving mouse retinal microvasculature and characterize mouse eye induced aberrations, we performed *in vivo* 2PFM angiography by retro-orbitally injecting dextran-conjugated fluorescein isothiocyanate (FITC) into the non-imaged eye. Aberrations were measured with fluorescence emitted from vessels in the superficial plexus (red asterisk, **Fig. 2A**; wavefront sensing area: 19 × 19 µm^2^). After AO correction, we observed a 2-10× enhancement in signal (**Fig. 2B,C**). Comparing the line signal profiles (along the orange dashed lines, **Fig. 2A,B**), we found that AO improved signal for all vessels while its impact on signal of smaller capillaries (**Fig. 2C**, black asterisks; 6-10× improvement) was more substantial than on larger vessels (**Fig. 2C**, black circles; 2-3× improvement). Despite the substantial signal improvements enabled by AO, we found that most capillaries, due to their size and sparse distribution in space, could be resolved in 3D without AO by our optimized 2PFM, albeit at reduced contrast and resolution (**Fig. 2D,E**). Our results indicate that a properly designed 2PFM is capable of acquiring retinal angiograms at the level of individual capillaries.

**Figure 2.**
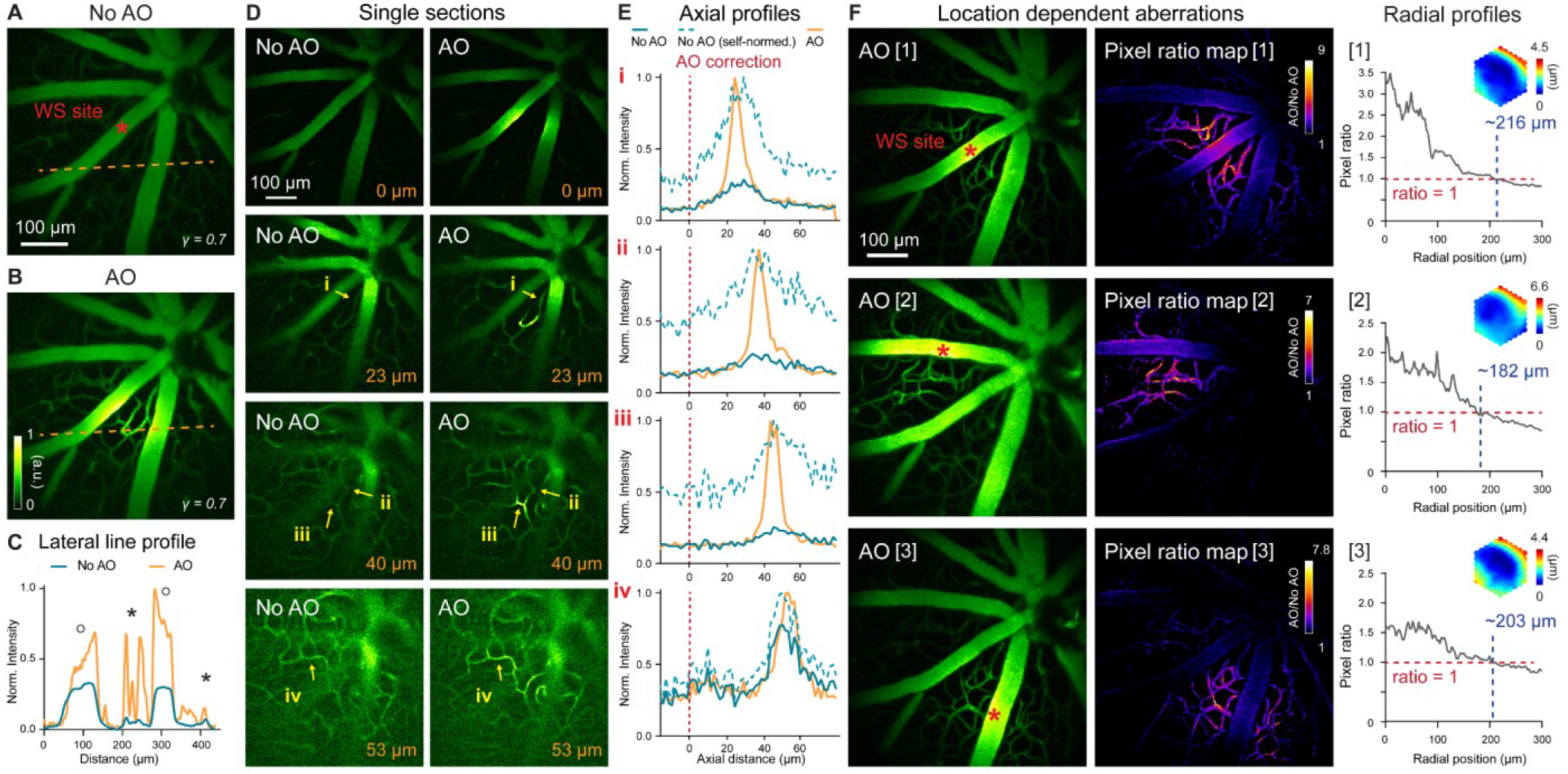
In vivo imaging of mouse retinal vasculature with AO-2PFM. **(A**,**B)** MIPs of image stacks (580 × 580 × 128 µm3) of vasculature measured (A) without and (B) with AO, respectively, normalized to AO image. Red asterisk: center of 19 × 19 µm2 wavefront sensing (WS) area. Gamma correction: 0.7. Representative data from > 25 experiments (technical replicates). **(C)** Lateral line profiles along orange dashed lines in A and B. Black circles: large vessels; black asterisks: capillaries. **(D)** Single image planes at 0, 23, 40, and 53 µm below the superficial vascular plexus acquired without and with AO correction performed at the superficial plexus (0 µm), normalized to AO images. **(E)** Axial profiles of capillary structures (i-iv in D). Red dashed lines: depth of wavefront sensing area. **(F)** Left: MIPs of image stacks (580 × 580 × 110 µm3) acquired with WS performed at different locations in the FOV (red asterisks). Middle: AO/No AO pixel ratio maps. Right: radially averaged profiles of pixel ratio maps, centered at WS sites. Insets: corrective wavefronts. MIPs and pixel ratio maps individually normalized.

We further evaluated how the mouse ocular aberrations varied with imaging depth and field position. We found that AO performed at the superficial plexus was beneficial for imaging deeper layers, with the correction at superficial depth improving signal, resolution, and contrast of deeper vasculature (**Figs. 2D,E**). This result indicated that most aberrations of the mouse eye arose from cornea and crystalline lens, instead of retina. Because the crystalline lens of the mouse eye has a gradient refractive index distribution^42,43^, ocular aberrations should also be field dependent^44,45^. Field-dependent aberrations might also be introduced when the mouse eye was positioned off-axis with respect to the eye imaging module. We therefore examined how aberrations varied with FOV position and characterized the area within which a single correction led to substantial signal improvement. We performed AO at different locations of the superficial plexus in the FOV (**Fig. 2F**, left column, red asterisks; Supplementary Movie 1) and compared their performance. The “AO/No AO” pixel ratio maps (**Fig. 2F**, middle column) exhibited field-dependent signal increase with larger gain achieved at pixels closer to the locations of aberration measurements. We quantified the effective area of AO in terms of signal improvement by calculating the radially averaged profiles of these pixel ratio maps (**Fig. 2F**, right column; origins at the wavefront sensing locations). We found signal improvement (“AO/No AO” pixel ratio ≥ 1) within a radius of ~216 µm when AO was performed at the FOV center of this mouse (**Fig. 2F**, [1]). For off-center locations, this radius was slightly smaller (**Fig. 2F**, [2] and [3]).

### AO enables 3D cellular resolution imaging of neurons in the mouse retina

The mouse retina consists of multiple layers of neurons with different cell types and distinct physiological properties. In the early stage of retinal diseases, abnormal morphology and function are usually confined to specific cell types within a single layer^46^. Therefore, for microscopic investigations of retinal physiology and pathology, it is essential to resolve cells in 3D. We evaluated whether our optimized 2PFM was capable of 3D cellular resolution imaging without correcting the severe aberrations of the mouse eye.

For this purpose, we imaged the densely fluorescent Thy1-YFP-16 mouse retina *in vivo*, where all bipolar cells, amacrine cells, and retinal ganglion cells were labeled with yellow fluorescence protein^34^ (YFP). A single AO correction acquired by scanning a 19 × 19 µm^2^ area (centered on the red asterisk in **Fig. 3A**) substantially improved signal and resolution (**Fig. 3A,B**; Supplementary Movie 2), recovering higher spatial frequency information in both lateral and axial images (**Fig. 3C,D**). The resolution enhancement was especially striking along the axial direction, allowing retinal layers to be more clearly differentiated by better resolving neurons at different depths (**Fig. 3A,B**, XZ images). This improvement in axial resolution is especially important for functional imaging, because it minimizes neuropil contamination and ensures accurate characterization of the functional properties of neurons^47–49^. Therefore, AO was necessary for 3D cellular resolution imaging of retinal neurons *in vivo*. In the lateral image planes, our optimized 2PFM design and mouse preparation allowed the identification of individual neurons without AO, albeit at lower signal and poorer resolution than those achieved with AO, for inner nuclear layer, inner plexiform layer, and ganglion cell layer (**Fig. 3E**). In contrast, subcellular processes could not be visualized without aberration correction (e.g., processes in the inner plexiform layer, **Fig. 3E**, middle column).

**Figure 3.**
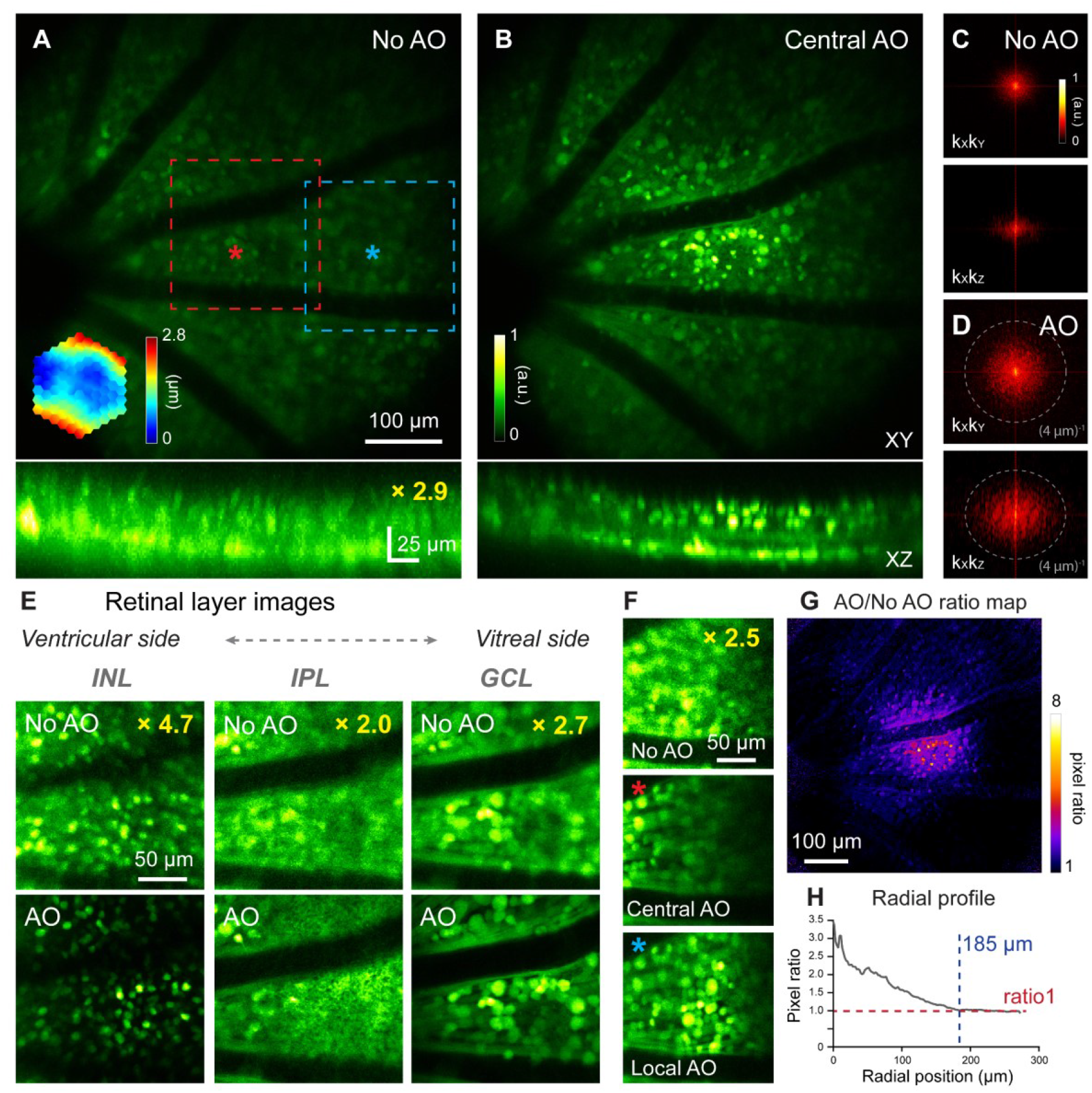
*In vivo* imaging of mouse retinal neurons with AO-2PFM. **(A**,**B)** MIPs of image stacks (580 × 580 × 80 µm3) of a Thy1-YFP-16 retina, measured (A) without and (B) with AO, respectively, normalized to AO images. Red asterisk: center of a 19 × 19 µm2 WS area. Top: lateral (XY) MIPs. Bottom: axial (XZ) MIPs; ‘No AO’ image brightness artificially increased by 2.9× for visualization. Representative data from > 10 experiments (technical replicates). **(C**,**D)** kXkY and kXkZ spatial frequency representation of images in (A,B). **(E)** Images of different retinal layers within the red dashed box in A acquired (top) without and (bottom) with AO, respectively, normalized to AO images. INL: inner nuclear layer; IPL: inner plexiform layer; GCL: ganglion cell layer. INL/GCL: MIPs of 4.9/7.8-µm-thick image stacks; IPL: single image plane. ‘No AO’ image brightness artificially increased for visualization (gains shown in each image). **(F)** Single image planes in GCL at FOV edge (blue dashed box in A) acquired (top) without AO, (middle) with central AO (WS area centered at red asterisk in A), and (bottom) with local AO (WS area centered at blue asterisk in A), respectively. Images normalized to local AO image. ‘No AO’ image brightness artificially increased by 2.5× for visualization. **(G)** AO/No AO pixel ratio map. **(H)** Radially averaged profile of pixel ratio map, centered at red asterisk in A.

Similar to our vascular imaging results, the Thy1-YFP-16 mouse eye exhibited field-dependent aberrations. For areas away from the AO measurement location (e.g., blue dashed box in **Fig. 3A**), although resolution improvement remained, the correction acquired at the FOV center (**Fig. 3F**, Central AO) did not increase signal strength as much as the locally acquired correction (centered on the blue asterisk in **Fig. 3A**; **Fig. 3F**, Local AO). For the Thy1-YFP-16 mouse, the effective area of AO performed at the FOV center was estimated from the “AO/No AO” ratio map (**Fig. 3G**) to have a radius of ~185 µm (**Fig. 3H**).

### Strategy for enlarging the effective area of AO correction for 3D cellular resolution imaging

Imaging retinal vascular and neuronal structures, we found that the spatially-varying aberrations of the mouse eye limited the effective area for AO correction that was acquired by sensing wavefront from a small region of the retina (e.g., 19 × 19 µm^2^ for **Figs. 1-3**). Although this approach succeeded in resolving synaptic features (**Fig. 1C**) and neuronal processes (**Fig. 3E,F**), for applications requiring 3D neuronal population imaging, synaptic resolution can be sacrificed in favor of cellular resolution imaging capability over larger FOVs. The latter can be achieved by correcting only for global mouse eye aberrations measured by scanning a larger retinal region for wavefront sensing.

As a demonstration, for a 580 × 580 µm^2^ FOV, we measured aberrations from areas of 19 × 19, 95 × 95, 190 × 190, and 380 × 380 µm^2^ (**Fig. 4A, i-iv**, yellow dashed boxes) and obtained differing corrective wavefronts resulting from the spatially varying aberrations. Quantifying and comparing AO effectiveness by their “AO/No AO” pixel ratio maps (**Fig. 4A**), we found that correcting aberrations from smaller areas provided greater local signal improvement but exhibited faster decay in signal improvement over distance (**Fig. 4B, i, ii**). This was because the corrective wavefront acquired from a small FOV completely cancelled out the local aberrations and led to diffraction-limited imaging of local structures. For structures away from the wavefront sensing region and thus experiencing different aberrations, however, the same corrective wavefront led to substantial residual aberrations that degraded AO performance. In contrast, correcting aberrations from a larger area reduced signal improvement in the center of the area but enlarged the overall area within which signal was enhanced, which now extended over the entire imaging FOV (**Fig. 4B, iii, iv**). Here, the wavefront measured from scanning the guide star over a larger FOV averaged out the local variations and represented the wavefront distortions common to all field positions. As a result, even though the improvement at the center of the wavefront sensing area was not as large, by removing the common aberrations from the entire FOV, this approach led to a larger effective area for AO correction.

**Figure 4.**
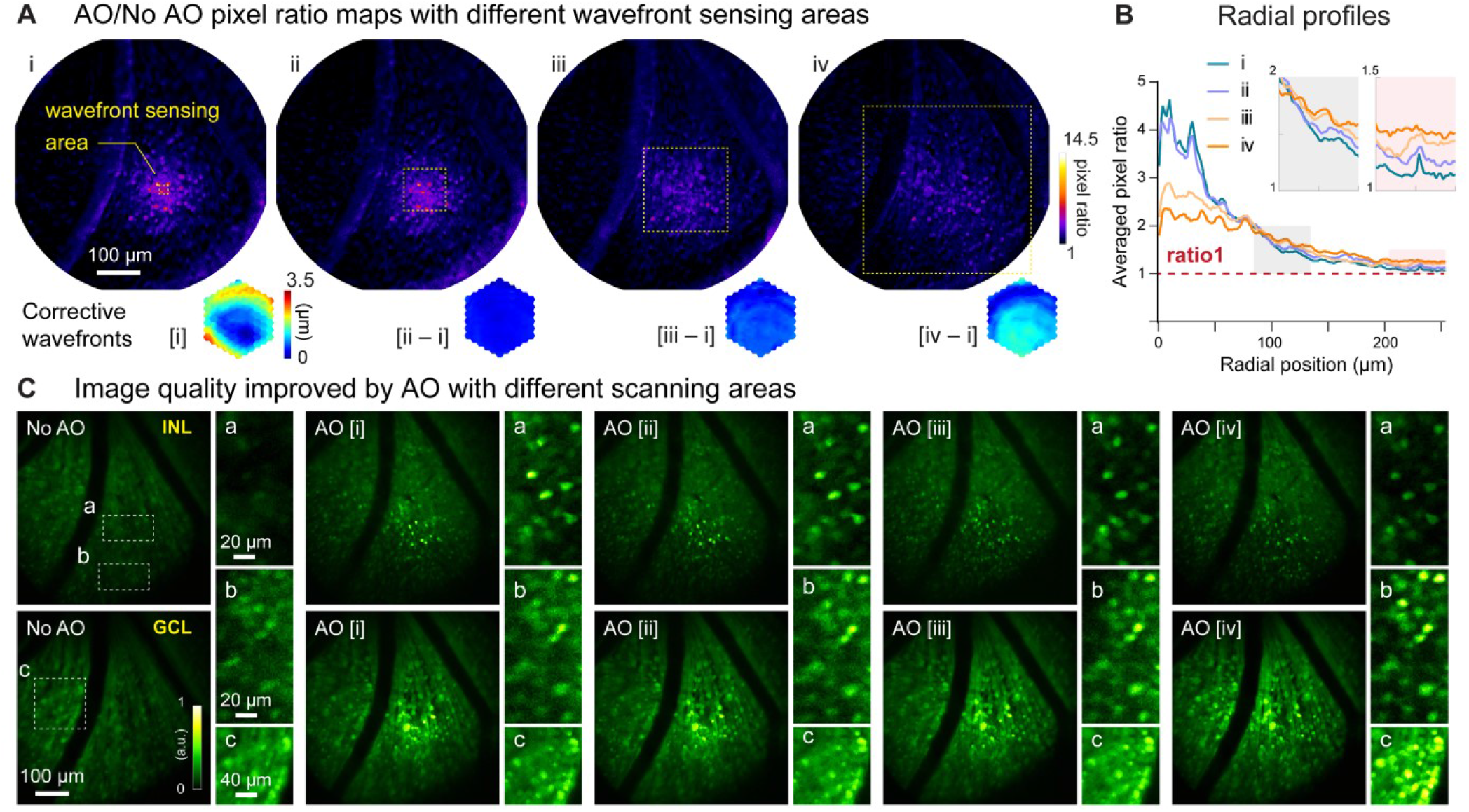
Larger WS areas enlarges the effective region of AO correction for 3D cellular resolution imaging. **(A)** Top: AO/NoAO pixel ratio maps for corrections with differently sized WS areas (yellow dashed boxes; i, 19 × 19 µm2; ii, 95 × 95 µm^2^; iii, 190 × 190 µm^2^; iv, 380 × 380 µm^2^). Bottom: (for [i]) corrective wavefront and (for [ii-iv]) difference in wavefronts between [ii-iv] corrective wavefronts and [i] corrective wavefront. **(B)** Radially averaged profiles of pixel ratio maps in (A). Insets: zoomed-in views of shaded areas. **(C)** Single image planes acquired (top) from INL and (bottom) GCL without and with AO using corrective wavefronts [i-iv], respectively. Insets: zoomed-in views of areas at FOV (a) center and (b,c) edge. All images normalized to AO images (AO [i] for inset a; AO [iv] for inset b,c).

Importantly, this approach enabled large-scale imaging of the retina with 3D cellular resolution, as indicated by retinal cell images taken from the center and edge locations (**Fig. 4C**). A more localized wavefront correction (e.g., AO [i], **Fig. 4C**) gave rise to sharper images at the scanning center (**Fig. 4C**, insets a), while a more global wavefront measurement (e.g., AO [iv], **Fig. 4C**) benefited more the visualization of neurons towards the edge of the FOV (**Fig. 4C**, insets b and c). Moreover, with global corrections, neuronal images at the center of the area maintained cellular resolution despite reduction in signal gain (**Fig. 4C**, insets a). Our results suggest that for diffraction-limited imaging of fine structures within a small FOV, a localized wavefront measurement is required, whereas a global wavefront measurement is preferable for 3D cellular resolution imaging over large FOVs.

### High-resolution *in vivo* identification of abnormal capillaries in a pathological mouse model

Having demonstrated the effectiveness of our AO-2PFM in improving signal, contrast, and spatial resolution for *in vivo* retinal imaging, we utilized our system to study retinal microvascular pathology. Retinal angiomatous proliferation (RAP), a subtype of age-related macular degeneration, is characterized by capillary proliferation that originates from the sensory retina and extends into the subretinal space^50^. Replicating the characteristic phenotypes of human RAP, a transgenic mouse model, the very low-density lipoprotein receptor knockout (VLDLR-KO) mouse, has been employed to study the underlying mechanism of RAP. In this model, the gene encoding VLDLR, which mediates anti-angiogenic signaling in retinal vasculature, is knocked out, leading to overgrown intraretinal vasculature and subretinal neovascularization^51,52^. In addition, fluorescein angiography revealed that the VLDLR-KO model of proliferative vascular retinopathy has extensive focal vascular leakage^51–54^. However, the lack of sufficient spatial resolution and optical sectioning capability makes it challenging for fluorescence angiography to identify the 3D location and characterize the structure of the vascular lesions *in vivo*.

We utilized AO-2PFM to image *in vivo* the retina of VLDLR-KO/Sca1-GFP and their wildtype control WT/Sca1-GFP mice, both with vascular endothelial cells in the retina labeled with GFP^54^. In order to detect microscopic capillary pathology, we used 19 × 19 µm^2^ wavefront sensing area to achieve diffraction-limited imaging performance, which led to high-resolution images of endothelial cell linings of retinal vessels in both mouse lines (**Figs. 5A,B**). Interestingly, in the VLDLR-KO/Sca1-GFP retina, images acquired with AO revealed a disruption in the capillary endothelium by Sca1-GFP labeling (**Fig. 5A**, yellow asterisks, insets i-ii; Supplementary Movie 3), which were not observed in the WT/Sca1-GFP retina (**Fig. 5B**). We further confirmed the presence of such microvascular lesions using *ex vivo* 2PFM imaging of dissected VLDLR-KO/Sca1-GFP retinas (**Fig. S3A,C;** Supplementary Movie 4). Whereas similarly structured capillary disruptions were observed in the VLDLR-KO/Sca1-GFP retina, consistent with the *in vivo* investigation, capillaries in the wildtype control had normal structures (**Fig. S3B,D;** Supplementary Movie 5).

**Figure 5.**
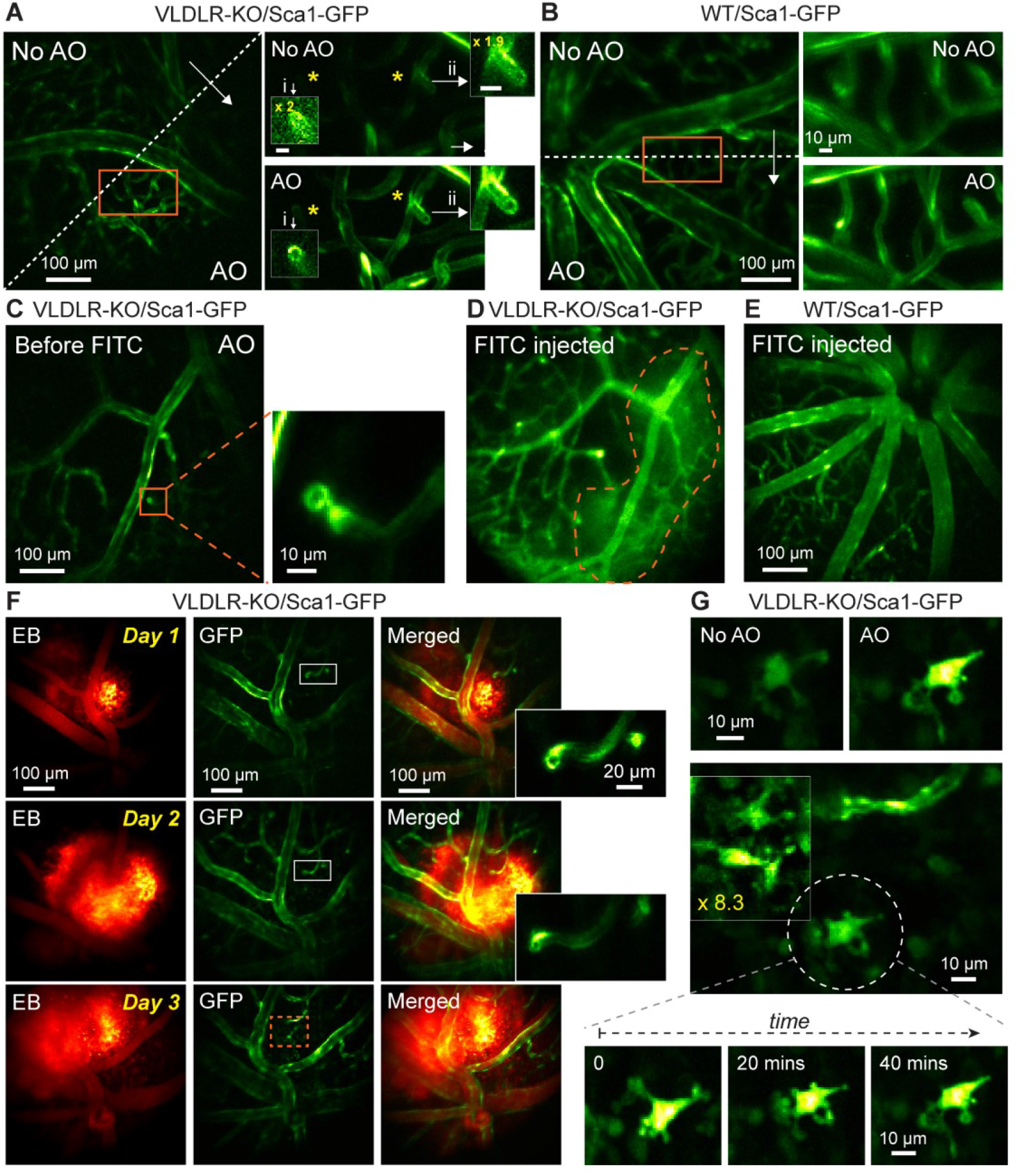
*In vivo* vasculature imaging in pathological and healthy retinas. **(A**,**B)** Left: MIPs of image stacks of (A) VLDLR-KO/Sca1-GFP (580 × 580 × 94 µm3) and (B) WT/Sca1-GFP (520 × 520 × 120 µm3) mouse retinas, measured (arrow start) without and (arrow end) with AO. Asterisks: capillary disruptions. Insets: zoomed-in views individually normalized for better visualization. ‘No AO’ inset brightness artificially increased for visualization (gains shown in inset). **(C)** A single image plane of a VLDLR-KO/Sca1-GFP mouse retina before FITC injection. Inset: MIP of a zoomed-in image stack (58 × 58 × 8.2 µm3) showing capillary lesion (orange box). **(D)** The same FOV in (C) after FITC injection. Dashed region: area with heightened fluorescence outside the vasculature. **(E)** MIP of an image stack (580 × 580 × 150 µm3) of a WT/Sca1-GFP mouse retina after FITC injection. **(F)** Retinal images taken on (top) day 1, (middle) day 2, and (bottom) day 3 after Evans Blue (EB) injection. Left: near-infrared channel showing EB-labeled vasculature and tissue staining (MIP of a 580 × 580 × 166 µm3 volume). Middle: green channel showing GFP-labeled vasculature (single planes). Right: merged images. Insets: zoomed-in views of gray rectangles from the GFP images (single sections). **(G)** Microglia observed in EB-injected VLDLR-KO/Sca1-GFP mouse retina on day 3 near the lesion site (orange dashed box in F). Top: microglia imaged without and with AO. Middle: multiple microglia in the leaking region. Signal in the boxed region was artificially increased by 8.3× for visualization. Bottom: time-lapse images of the microglia in white dashed circle. All images are single planes. Wavefront sensing area: 19 × 19 µm2. *In vivo* data in this Figure were obtained from 3 VLDLR-KO and 2 WT mice (biological replicates).

We hypothesized that these lesions as capillary disruptions observed in the VLDLR-KO/Sca1-GFP retina were the locations of dye leakage. To test this hypothesis, enabled by AO, we first located a microvascular lesion in a VLDLR-KO/Sca1-GFP mouse retina (**Fig. 5C**, orange box, inset). Then we retro-orbitally injected the green fluorescent dye FITC into the non-imaged eye, which labeled the blood plasma within the retinal vasculature (**Fig. 5D**). Immediately after dye injection, we observed dye leakage around the lesion site (**Fig. 5D**, orange dashed area). A control experiment was carried out by introducing FITC into the healthy WT/Sca1-GFP mouse retina retro-orbitally, where neither capillary disruptions nor dye leakage were observed (**Fig. 5E**).

To further study the association between dye leakage and microvascular lesions, we injected the NIR dye Evans Blue (EB) into the retinal vasculature and performed dual-color 2-photon imaging of the VLDLR-KO retina. Similar to the experiments with FITC, we observed leakage in the knockout mouse retina, with EB persistently staining retinal tissue and the stained volume expanding over three days of consecutive imaging (**Fig. 5F**). We observed capillary lesions (**Fig. 5F**, insets) in the stained volume, suggesting a spatial correlation between dye leakage and capillary abnormalities. Moreover, on the third day, we observed microglia within the dye-stained retinal volume that did not show up in previous two days (**Fig. 5G**), suggesting that the leakage of EB triggered local immune response and recruited microglia to the impacted area. In addition, with AO-2PFM, we were able to track morphological changes in the processes of the same microglia at subcellular resolution (**Fig. 5G**, bottom). Control experiment in WT/Sca1-GFP retina showed local small-scale EB leakage (**Fig. S4**), probably resulting from normal remodeling of the retinal vasculature^2^. Our findings revealed, for the first time, the microscopic morphological details of vasculature lesions and suggested that these capillary disruptions served as intraretinal origins of vascular leakage in the VLDLR knockout mouse. Here AO was essential for 2PFM to achieve high-resolution identification and characterization of microvasculature lesions *in vivo*. Together with our optimized sample preparation, AO-2PFM also allowed us to track these lesions, dye leakage, and associated immune response longitudinally, making it possible to investigate the development and progression of vasculature-associated diseases at subcellular resolution *in vivo*.

### High-resolution *in vivo* imaging of retinal pharmacology

With the 3D cellular resolution imaging capability enabled by AO-2PFM, we can now image the functional activity of retinal neurons with high fidelity in healthy or diseased retina *in vivo* using activity sensors such as the genetically encoded calcium indicator GCaMP6s^55^.

As a demonstration, we studied how pharmacological manipulation affects RGC activity *in vivo* in a mouse model of retinal degeneration. As the afferent neurons of the retina, RGCs deliver retinal circuit output to the rest of the brain and play a crucial role in visual perception. RGCs in the *rd1* mouse, the oldest and most widely studied animal model of retinal degeneration^56^, become hyperactive after photoreceptor death caused by a mutation in the *Pde6b* gene^57,58^. Recent studies have suggested that RGC hyperactivity masks light-evoked signals initiated by surviving photoreceptors and impedes remaining light-elicited behaviors^58,59^. Studying RGC hyperactivity therefore is of great importance both for understanding the pathology of retinal degeneration and for developing pharmacological therapies^60^. However, RGC hyperactivity has been only studied *ex vivo* on dissected retinas^58,60^, preventing longitudinal evaluation of degeneration progression and therapeutic approaches.

Here we characterized RGC hyperactivity *in vivo* and studied the effect of Lidocaine, a use-dependent Na^+^ channel blocker, on alleviating hyperactivity of RGCs in the *rd1*-Thy1-GCaMP6s mouse using AO-2PFM and calcium imaging. Because RGC hyperactivity is usually studied by *ex vivo* tools such as multi-electrode array (MEA) or single cell electrophysiology recordings, to establish the calcium signature of RGC hyperactivity, we first carried out simultaneous cell-attached and *ex vivo* 2PFM calcium recordings of the same hyperactive alpha RGCs in a dissected *rd1* mouse retina (**Fig. 6A**). Consistent with previous reports on *ex vivo* retina^58–60^, RGC hyperactivity was observed as high-frequency action potentials. In terms of calcium signaling (quantified as calcium response magnitude ΔF/F, with F being baseline brightness and ΔF being the difference from baseline brightness), hyperactivity measured *ex vivo* was correlated with a heightened ΔF/F of the GCaMP6s-expressing soma. A temporally varying firing rate led to fluctuations in the ΔF/F of its calcium signal. After ~20 seconds of 2% Lidocaine bath perfusion, spontaneous spiking from the RGC was largely suppressed with a ΔF/F close to 0. After artificial cerebrospinal fluid (ACSF) washout, RGC hyperactivity partially recovered, which was associated with an increase of ΔF/F magnitude. The observed time course and suppressive effect of Lidocaine application on RGC hyperactivity were consistent with *ex vivo* multi-electrode array (MEA) recordings (**Fig. S5A,B**). The characteristics of the corresponding calcium responses were also observed in 2-photon population imaging of multiple RGCs in dissected *rd1* retina (**Fig. 6B**), with the brightness and ΔF/F of the GCaMP6s-expressing neurons reduced by Lidocaine application and followed by partial or full recovery after washout.

**Figure 6.**
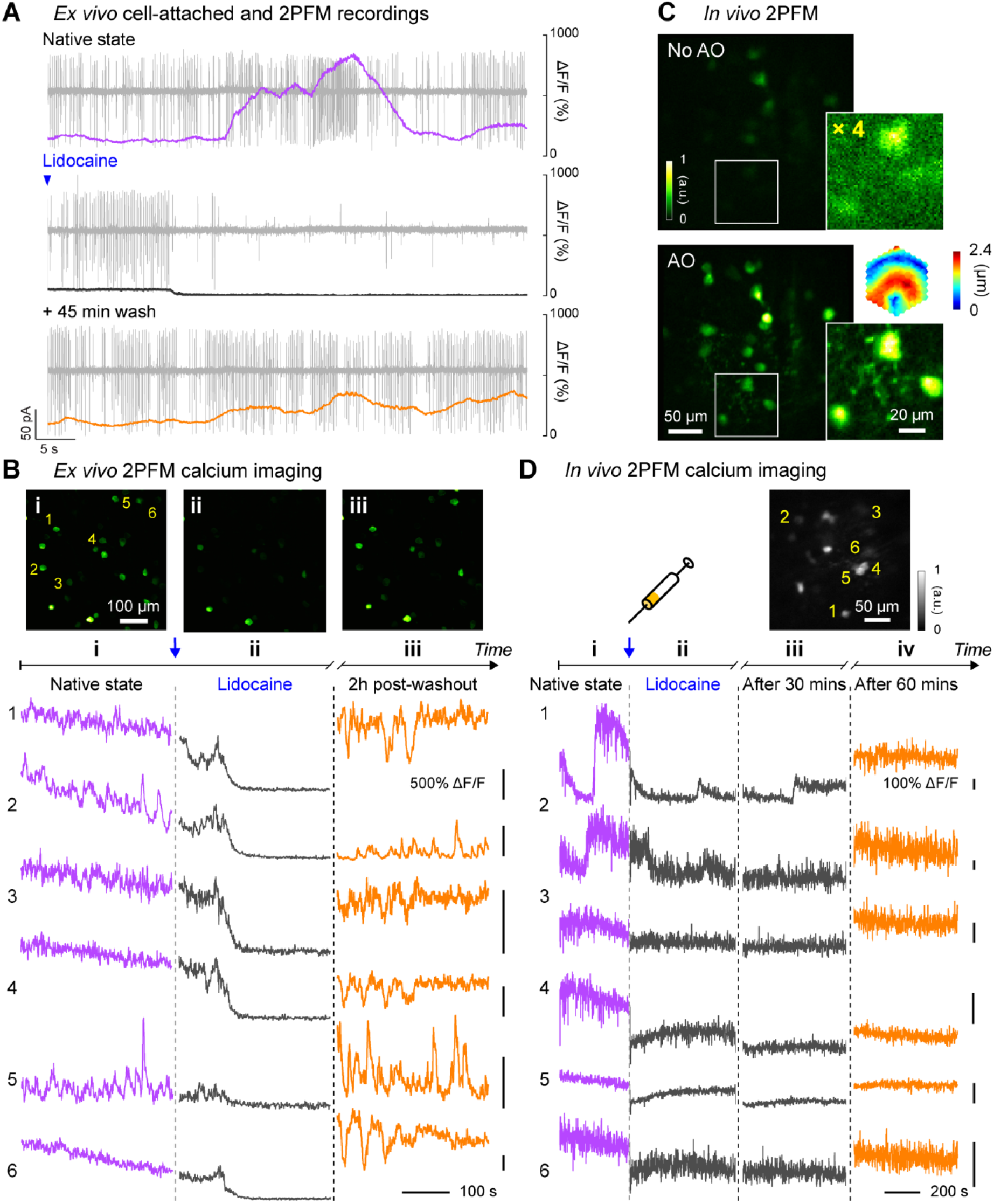
*In vivo* calcium imaging of Lidocaine-suppressed RGC hyperactivity in *rd1*-Thy1-GCaMP6s mouse retina. **(A)** Simultaneous cell-attached and 2PFM calcium recordings of a RGC before, during, and 45 mins after Lidocaine treatment. Representative data from > 3 cells. **(B)** Top: average intensity projections of *ex vivo* 2PFM images of RGCs in a dissected retina (i) before, (ii) right after, and (iii) 2 hours after Lidocaine treatment, normalized to the left image. Bottom: *Ex vivo* calcium dynamics of 6 RGCs therein. Representative data from > 3 retinas. (**C)** *In vivo* single image planes of RGCs acquired without and with AO, respectively, normalized to AO images. Insets: zoomed-in views and corrective wavefront; ‘No AO’ inset brightness artificially increased by 4.0× for visualization. Representative data from > 3 retinas. (**D)** *In vivo* calcium dynamics of 6 RGCs (i) before, (ii) right after, (iii) 30 minutes after, and (iv) 60 minutes after Lidocaine treatment, respectively. Wavefront sensing area: 19 × 19 µm2. Representative data from 1 retina.

Having confirmed that RGC hyperactivity was associated with heighted calcium levels, we next performed AO-2PFM calcium imaging to directly study how Lidocaine affected RGC hyperactivity *in vivo*. Through the *rd1*-Thy1-GCaMP6s mouse eye, AO increased RGC brightness by on average 4× and enabled high-resolution visualization of both RGC somata and their processes (**Fig. 6C**). The signal increase enabled by AO was particularly important for the *rd1*-Thy1-GCaMP6s mouse, because the RGCs here had dimmer fluorescence than the other lines that we investigated. For these RGCs, correcting the eye-induced aberration was essential for their visualization and high-fidelity functional investigations at cellular resolution *in vivo*. To maximize the fluorescence signal, we performed AO with a small (19 × 19 µm^2^) wavefront sensing area. Before injecting Lidocaine, we observed slow fluctuations in the brightness of GCaMP6s-expressing RGCs (**Fig. 6D, i**), similar to the slow dynamic events in RGC calcium traces measured *ex vivo*. One minute after retro-orbital injection of Lidocaine into the non-imaged eye, hyperactivities from these cells were substantially inhibited for an hour as indicated by the reduction of RGC GCaMP6s fluorescence brightness (**Fig. 6D, ii and iii**). Imaging the same RGCs 60 mins after injection (**Fig. 6D, iv**), we detected partial to complete recovery of RGC brightness, consistent with our *ex vivo* recordings after washing out. Here, by studying the suppression effects of Lidocaine on RGC hyperactivity within living mice, we demonstrated that AO-2PFM can monitor the pathology and pharmacology of retinal diseases at high resolution *in vivo*.

## Discussion

By optimizing optical design and sample preparation and using direct wavefront sensing AO to correct mouse eye aberrations, we demonstrated here the first *in vivo* visualization of retinal synaptic structures, the first *in vivo* identification of capillary lesions with sub-capillary details, and the first *in vivo* detection of RGC hyperactivity and its suppression by pharmacological reagents.

To image mouse retina *in vivo* with 2PFM, one can either utilize a standard objective lens^19,40^ or the mouse eye’s optics itself^23,26^ to focus the excitation light and collect the fluorescence emission. The former approach requires long-working-distance objective lenses and, more importantly, suffers from severe aberrations caused by the refractive power of the ocular optics (mostly crystalline lens, as cornea was typically flattened in these systems). For this reason, to achieve the best image quality and the largest FOV size, using mouse eye itself as the focusing element as implemented here is preferable.

Among the studies that used the mouse eye optics for imaging, discrepancies exist in how large the mouse ocular aberrations are and how essential AO is for vasculature and cellular imaging in the mouse retina. For the multiple mouse strains investigated here (i.e., wild-type (C57BL/6), Thy1-GFP line M, Thy1-YFP-16, VLDLR-KO/Sca1-GFP, WT/Sca1-GFP, and *rd1*-Thy1-GCaMP6s), we found that their ocular aberrations were typically within the range of 3~5 µm peak-to-valley (P-V; after removing tip, tilt, and defocus) and 0.4~0.8 µm rms (**Fig. S6**) without notable differences in severity across strains. While consistent with most previously reported values^21,25^, our study differs significantly from a recent AO-2PFM study that reported extremely large aberrations (e.g., 12-25 µm P-V^26^) and found it difficult to resolve microvasculature and cell bodies in 2D *in vivo* without AO. In contrast, with our carefully designed microscope and system aberration correction procedure, we achieved capillary visualization, 2D single-cell resolution, and retinal layer differentiation by only correcting system aberrations (‘No AO’ in our case). Given that our system aberrations were much smaller than mouse ocular aberrations (**Fig. S6**), our study indicated that a well-engineered 2PFM like ours should be sufficient for *in vivo* retinal imaging applications requiring only capillary and 2D cellular resolution.

One factor that strongly impacted image and wavefront sensing quality was the sample preparation procedure. We found that our specially designed 0-diopter contact lens encircled with a supportive flat base were essential for high-quality imaging by 2PFM both without and with AO (**Fig. S2**). Similar improvement in imaging quality by 0-diopter contact lens was reported previously for *in vivo* optical coherence tomography imaging of the rat retina, where it was hypothesized that the application of the contact lens smoothed corneal defects and reduced wavefront error of the anterior segment of the eye^61^. The supportive flat base encircling the optical zone of our contact lens^62^ and the gel completely separated the eye surface from air and prevented cataract formation. Together, they enabled high-quality SH images and accurate corrective wavefronts to be acquired throughout the experiment.

Incorporating direct-wavefront-sensing-based AO with 2PFM, we found that location-dependent aberrations led to local improvement in the mouse retina *in vivo*. To enlarge the high-resolution area enabled by AO, one way is to stitch images from smaller areas, each with its own local AO correction^44,45,63^. However, this procedure can be time-consuming and thus nonideal for *in vivo* functional studies. By scanning differently sized areas for wavefront sensing, we identified a trade-off between AO performance (i.e., resolution and signal enhancement) and effective area. We demonstrated that a single corrective wavefront acquired by scanning the guide star over a more extended area led to 3D cellular resolution imaging over a larger retinal volume, simplifying the procedure for future functional studies of neuronal populations in the retina. It is worth noting that, instead of incorporating scan lenses optimized for large scanning angles, our homebuilt 2PFM system utilized regular achromatic doublet lenses, which reduced cost but limited the overall imaging FOV of our microscope. The effective AO area and imaging FOV would be further increased by incorporating high-performance scan lenses.

To study retinal pathologies for physiological and clinical insights, it is ideal to conduct longitudinal investigations *in vivo*. Importantly, to probe microscopic early-stage pathologies, high spatial resolution is needed. In human applications, investigation of retinal vascular abnormalities are limited to capillary resolution^64,65^. Sub-capillary morphology and dynamics of the mouse retina were recently observed by light-sheet microscopy, however with *ex vivo* preparations^66^. To our knowledge, sub-capillary features had not been observed in the living mouse eye previously. In this work, applying AO-2PFM, we studied retinal vasculature in a pathological mouse model with proliferative vascular retinopathy at sub-capillary resolution *in vivo*. Recovering diffraction-limited resolution, AO enabled us to identify capillary lesions as capillary endothelium disruptions that were associated with dye leakage in 2-photon fluorescence angiograms. Moreover, the repeatable and reliable AO performance allowed us to track the same retinal region over multiple days and discover lesion-associated immune response and microglia migration at subcellular resolution. Our results show far-reaching potential of AO-2PFM for mechanistic understanding and early diagnosis of retinal diseases.

We also applied our AO-2PFM to *in vivo* activity imaging of RGCs in a mouse model of retinal degeneration. Due to the dimmer brightness of the fluorescence indicator in this model, AO was essential in increasing signal strength and enabling high-sensitivity interrogation of the effects of pharmacological manipulation on RGC hyperactivity. Traditionally, pharmacological effects on retina are studied by electrophysiological and imaging tools on *ex vivo* retinal preparation, or *in vivo* by indirect assessments downstream in the visual pathway or through behavior test^59^. Taking retinal degeneration as an example, treatment-induced photosensitization enhancement has been mainly evaluated through electrophysiology or *ex vivo* imaging of dissected retinas. AO-2PFM enabled us to evaluate how pathological RGC hyperactivity was suppressed by an example pharmacological agent, lidocaine, at single cell level noninvasively. Together with the capability for longitudinal investigations discussed above, we envision that the AO-enabled high-sensitivity subcellular and cellular 2PFM imaging would become a highly enabling technology for pathological and pharmacological investigations of the mouse retina *in vivo*.

## Materials and methods

### Animal use

All animal experiments were conducted according to the National Institutes of Health guidelines for animal research. Procedures and protocols (AUP-2020-06-13343) were approved by the Institutional Animal Care and Use Committee at the University of California, Berkeley.

### AO two-photon fluorescence microscope (AO-2PFM)

The AO-2PFM was built upon a homebuilt 2PFM (**Fig. 1A**) incorporated with a direct-wavefront-sensing-based AO module, as described in detail previously^27^. Briefly, 920-nm output from a femtosecond Ti:Sapphire laser (Coherent, Chameleon Ultra II) was expanded (2×, Thorlabs, GBE02-B) after a Pockel Cell (ConOptics, 350-80-LA-02-BK). The beam was then scanned with a pair of optically conjugated (by L1-L2, FL = 85 mm; Edmund Optics, 49-359-INK) galvanometer mirrors (Cambridge Technology, 6215H). A pair of achromatic lenses (L3-L4, FL = 85 and 300 mm; Edmund Optics, 49-359-INK and 49-368-INK) relayed the galvos to the DM (Iris AO, PTT489). The focal plane position of two-photon excitation in the mouse retina was controlled by an electrically tunable lens (ETL; Optotune, EL-16-40-TC-VIS-5D-C), which was conjugated to the DM (by L5-L6, FL = 175 and 400 mm; Edmund Optics, 49-363-INK and Newport, PAC090). The ETL was then relayed to the pupil of the mouse eye by L7 (FL = 200; Thorlabs, AC254-200-AB) and L8, which was composed of two identical lenses (FL = 50 mm; Thorlabs, AC254-050-AB). The two 50-mm-FL lenses in L8 were used together with a combined FL of 25 mm, and they were mounted with their curved surfaces facing and almost touching each other (**Fig. S1A,D**) to minimize aberrations during large-angle scanning. For 2PFM imaging, the emitted fluorescence from the mouse retina was collected by the mouse eye, travelled through L8-L7 and the ETL, reflected by a dichroic mirror (D2; Semrock, Di02-R785-25×36), focused by a lens (L9, FL = 75 mm; Thorlabs, LB1309-A), and detected by a photomultiplier tube (PMT, Hamamatsu, H7422-40). For direct wavefront sensing, D2 was moved out of the light path and the emitted fluorescence was descanned by the galvo pair, reflected by a dichroic mirror (D1; Semrock, Di03-R785-t3-25×36), and relayed to a Shack-Hartmann (SH) sensor by a pair of lenses (L10-L11, FL = 60 and 175 mm; Edmund Optics, 47-638-INK and 47-644-INK). The SH sensor was composed of a lenslet array (Advanced Microoptic System GmbH, APH-Q-P500-R21.1) and a camera (Hamamatsu, Orca Flash 4.0) that was placed at the focal plane of the lenslet array. Wavefront aberrations were measured from the shifts of SH pattern foci, reconstructed with custom MATLAB code, and the corresponding corrective pattern was then applied to the DM.

### System correction

Before imaging the mouse retina, system aberration caused by imperfect and/or misaligned optics was corrected. Due to the path difference between the two-photon illumination and the fluorescence wavefront sensing paths^67^, system correction was performed with a modal-based optimization approach^25,68^. Specifically, with 0 mA applied to the ETL, we imaged a fluorescent lens tissue sample at the focal plane of L7 and applied 11 values (−0.1 ~ 0.1 µm rms at an increment of 0.02 µm) for each of the first 21 Zernike modes excluding piston, tip, tilt, and defocus. The optimal value for each Zernike mode was determined by maximizing the fluorescence intensity of the sample and it was applied to the DM before proceeding to the next Zernike mode. An SH pattern was obtained with system aberration corrected and was used as the SH reference for calculating sample-induced aberrations. All images taken with system correction were indicated in the main text as “No AO”.

To change the focal plane within the retina, we varied the electric current applied to the ETL. We characterized how system aberrations varied with the ETL current (**Fig. S1**). We carried out system correction with 0 mA ETL current applied (**Fig. S6A**). Additional aberrations introduced by setting ETL current to 20, 40, 60, and 80 mA were negligible (**Fig. S1B**) compared with eye-induced aberrations (**Fig. S6**) and minimally affected *in vivo* imaging (**Fig. S1C**). We also evaluated how system aberrations varied with the distance D between the mouse eye pupil and the imaging module (**Fig. S1D**). Using Zemax® for ray tracing, we found its effect to be similarly minimal (**Fig. S1E**). Our typical *in vivo* retinal imaging was performed with 10~60 mA of ETL currents and 2~4 mm D values (**Fig. S1D**). Simulating the mouse eye as an ideal lens behind a 0-diopter contact lens (**Fig. S2A**) made of PMMA (1.49 refractive index) and 0.5-mm-thick eye gel (1.33 refractive index), we calculated the focal shifts and FOVs for different ETL currents using Zemax® and found a linear focal shift with ETL current and relatively constant FOV during 3D imaging (**Fig. S1F**). Imaging FOV and axial shift were determined from Zemax® simulation for D = 2 mm.

### *In vivo* imaging

All mice (Wild-type C57BL/6J and Thy1-YFP-16, the Jackson laboratory; VLDLR-KO/Sca1-GFP and WT/Sca1-GFP, Gong lab; GCaMP6s-*rd1*, Kramer lab) were at least 8 weeks old at the time of imaging. *In vivo* imaging was carried out on mice under isoflurane anesthesia (~ 1.0% by volume in O2). Prior to imaging, the mouse pupil was dilated with one drop of 2.5% phenylephrine hydrochloride (Paragon BioTeck, Inc) and one drop of 1% tropicamide (Akorn, Inc). A 0-diopter customized rigid contact lens (**Fig. S2A**, Advanced Vision Technologies) was placed on the eye, with eye gel (Genteal) applied in between the eye and the contact lens to prevent cornea drying and clouding. Excessive eye gel was removed by gently pressing the contact lens onto the mouse eyeball. One single application of eye gel was sufficient in keeping the cornea moist for a 2~4-hour imaging session. During imaging, mice were stabilized on a bite-bar on a 3D translational stage with two rotational degrees of freedom (Thorlabs, PR01) and the body temperature was maintained with a heating pad (Kent Scientific, RT-0515). The mouse head was carefully aligned to make the eye perpendicular to the illumination beam, minimizing off-axis aberrations and illumination clipping by the contact lens and mouse pupil. Fluorescent dyes were injected retro-orbitally into the non-imaged eye. In wild-type mice, 40-80 µL of 5% (w/v) 2M-Da dextran-conjugated FITC was injected for vasculature visualization. In some VLDLR-KO/Sca1-GFP mice, 30-40 µL of 5 mg/mL FITC or 5 mg/mL Evans Blue were injected. To generate bright enough fluorescent guide star for direct wavefront sensing in the weakly-fluorescent mouse line *rd1*-Thy1-GCaMP6s, 20-40 µL of 5 mg/mL Evans Blue was injected. To suppress RGC hyperactivity in *rd1*-Thy1-GCaMP6s mouse retina, we retro-orbitally injected 10 µL of 2% Lidocaine into the non-imaged eye.

All imaging parameters, including laser power at the mouse pupil, are listed in Table S1.

### Retina dissection

Mice were first euthanized by isoflurane overdose followed by cervical dislocation. Then the eyes were removed, and the retinas were isolated and immersed in standard oxygenated (95% O2, 5% CO2) artificial cerebrospinal fluid (ACSF) at room temperature and pH 7.2.

### *Ex vivo* 2-photon structural imaging of dissected Sca1-GFP mouse retinas

A commercial 2-photon fluorescence microscope (Bergamo^®^, Thorlabs) was used to image dissected Sca1-GFP mouse retinas (**Fig. S3**). 2-photon excitation at 920 nm was provided by a femto-second laser (Coherent, Chameleon Ultra II). *Ex vivo* images were acquired by a 16×0.8NA water-dipping objective lens (Nikon). Hardware controls and data acquisition were performed by ThorImage.

### Multielectrode array (MEA) recordings

Isolated *ex vivo rd1*-Thy1-GCaMP6s retinas were cut into three pieces. Each piece was mounted onto a 60-electrode MEA chip (60ThinMEA200/300iR0ITO, Multichannel Systems) with the inner retina facing the array, so that RGCs were in close contact with electrodes. The chip was connected to an amplifier (MEA1060, Multichannel Systems) for wide-band extracellular recording of multi-unit activity. Before the onset of recording, the retina was perfused with oxygenated ACSF at 34 °C for 30 min with a flowrate of 1 mL/min.

For pharmacological blockade of actional potentials, Lidocaine (2% in saline) was applied to the bath during corresponding recordings. Washout of lidocaine was performed by continuously perfusing oxygenated ACSF at 34ºC over the course of two hours with a flowrate of 1 mL/min.

Recorded activity from RGCs were high-pass filtered at 200 Hz, digitized at 20 kHz, and analyzed offline. Extracellular spikes were defined as transient signals with peak deflection of >3.5 standard deviations from the root mean square of background signal. Because individual electrodes can detect spikes from multiple RGCs, we utilized principal component analysis to sort unique units (Offline Sorter v3, Plexon), which accepted units having interspike intervals >1 ms. Each unit was compiled into a raster plot. The analysis code for processing sorted spike data into rasters is available online (https://github.com/kookstance/Multielectrode-array).

### Cell attached recordings of alpha-RGCs in *rd1*-Thy1-GCaMP6s retinas

Isolated *ex vivo rd1*-Thy1-GCaMP6s retinas were mounted onto filter paper (0.45 mm nitrocellulose membranes, MF-Millipore) with an optical window with the ganglion cell layer facing up. RGCs were visualized with DODT contrast infrared optics (Luigs and Neumann) and were targeted for whole cell recording with glass electrodes (4-6 MOhm) filled with ACSF. Loose-(< 1 GΩ) and tight-seal patches (> 1 GΩ) were obtained under voltage clamp with the command voltage set to maintain an amplifier current of 0 pA. Input resistance and series resistance were monitored throughout recording to ensure stable recording quality and cell health.

### *Ex vivo* 2-photon calcium imaging of *rd1* mouse retina

2-photon calcium imaging of *rd1*-Thy1-GCaMP6s retina was carried out on a custom galvo-scanning microscope (https://wiki.janelia.org/wiki/display/shareddesigns/Non-MIMMS+in+vivo+microscope) equipped with a 20× 1.0 NA water immersion objective (XLUMPLFLN20XW, Olympus). Excitation at 920 nm was provided by a tunable Ti:Sapphire ultrafast laser (Chameleon Ultra, Coherent). Imaging parameters were controlled by ScanImage 3.8.1 software (http://scanimage.vidriotechnologies.com/): 256×256 pixels at 1.25 Hz (2 ms per line). GCaMP6s emission was collected with a GaAsP PMT shielded by a longpass filter (ET500lp, Chroma).

Isolated retinas were cut into four-leaf clovers and transferred onto filter paper (0.45 mm nitrocellulose membranes, MF-Millipore) with the ganglion cell layer facing up. Oxygenated ACSF was then perfused over the retina at 34ºC for 30 minutes with a flowrate of 1 mL/min. An initial imaging session performed to account for potential 2-photon sensitivity. Experimental imaging was performed with the laser power at the sample ≤ 5 mW. For pharmacological blockade of actional potentials, Lidocaine (2% in saline) was applied to the bath during corresponding recordings. Washout of lidocaine was performed by continuously perfusing oxygenated ACSF at 34ºC over the course of two hours with a flowrate of 1 mL/min.

### Image processing and analysis

All image processing, visualization, and analysis were performed in ImageJ^69^. To remove motion-induced artifacts, image registration (TurboReg and StackReg plugins) was performed.

## Acknowledgements

We thank Tyson Kim and Bingyao Tan for helpful discussions. This work was supported by US National Institutes of Health grants U01NS103489 (Q.Z., W.C., and N.J.), U01NS118300 (Q.Z. and N.J.), RF1MH120680 (Y.Y., W.C., and N.J.), F32EY029983 (K.J.C.), R01EY024334 and P30EY003176 (R.H.K.), NIH/EY013849 (S.P., C.H.X., and X.G.).

## Data availability

All data generated or analyzed in this study are included in the manuscript and supporting files. Source data files have been provided for all primary figures. The Zemax file of the eye imaging module has been provided.

## Supplementary Materials for

## Other Supplementary Materials for this manuscript include the following

<u>Supplementary Movie 1</u>

*In vivo* 2-photon image stacks of retinal vasculature in a wildtype mouse measured without and with AO performed at different locations in the field of view (red asterisks). Same data as shown in Fig. 2F. Image volume: 580 × 580 × 110 µm^3^; Z step: 3.26 µm.

<u>Supplementary Movie 2</u>

*In vivo* 2-photon image stacks of retinal neurons in a Thy1-YFP-16 mouse measured without and with AO. Same data as shown in Fig. 3A,B,E. Image volume: 580 × 580 × 80 µm^3^ / 193 × 193 × 50 µm^3^; Z step: 1.63 / 0.98 µm.

<u>Supplementary Movie 3</u>

*In vivo* 2-photon image stacks of abnormal retinal capillaries in a VLDLR-KO/Sca1-GFP mouse measured without and with AO. Same data as shown in Fig. 5A. Image volume: 48 × 48 × 65 µm^3^; Z step: 3.26 µm.

<u>Supplementary Movie 4</u>

*Ex vivo* 2-photon image stacks of abnormal capillaries in dissected VLDLR-KO/Sca1-GFP mouse retinas. Same data as shown in Fig. S3A,C. Image volume: 1380 × 1380 × 82 µm^3^ / 78 × 78 × 82 µm^3^; Z step: 0.5 µm.

<u>Supplementary Movie 5</u>

*Ex vivo* 2-photon image stacks of normal capillaries in dissected WT/Sca1-GFP mouse retinas. Same data as shown in Fig. S3B,D. Image volume: 1380 × 1380 × 71 µm^3^ /78 × 78 × 71 µm^3^; Z step: 0.5 µm.

**Figure S1.**
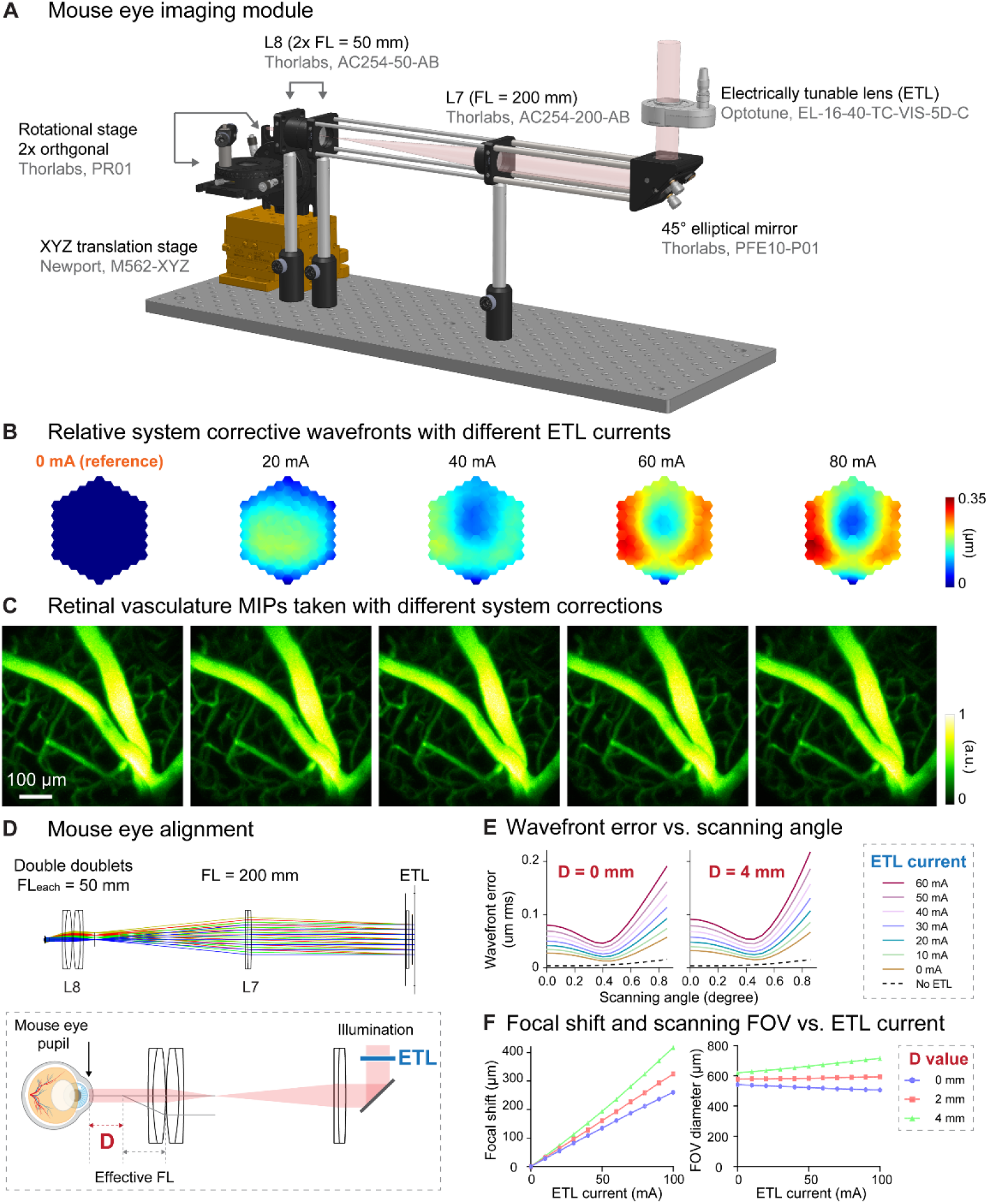
Characterization of aberrations introduced by ETL and alignment. **(A)** 3D rendering of the eye imaging module. **(B)** Additional corrective wavefronts for system aberrations measured with 0, 20, 40, 60, and 80 mA ETL currents, relative to the corrective wavefront measured with 0 ETL current. **(C)** MIPs of dye-injected retinal vasculature image stacks (550 × 550 × 127 µm3) measured with corresponding system corrections in (B). **(D)** Top: Zemax ray tracing of the eye imaging module. Bottom: illustration of mouse eye alignment (not to scale). D: distance between the mouse eye pupil and L8 focal plane. **(E)** Wavefront errors versus scanning angle (at the ETL) for different ETL currents with the mouse eye placed at D = 0 mm and D = 4 mm, respectively, obtained by ray tracing. **(F)** Relative focal shift and imaging FOV versus ETL current at different D values, obtained by ray tracing.

**Figure S2.**
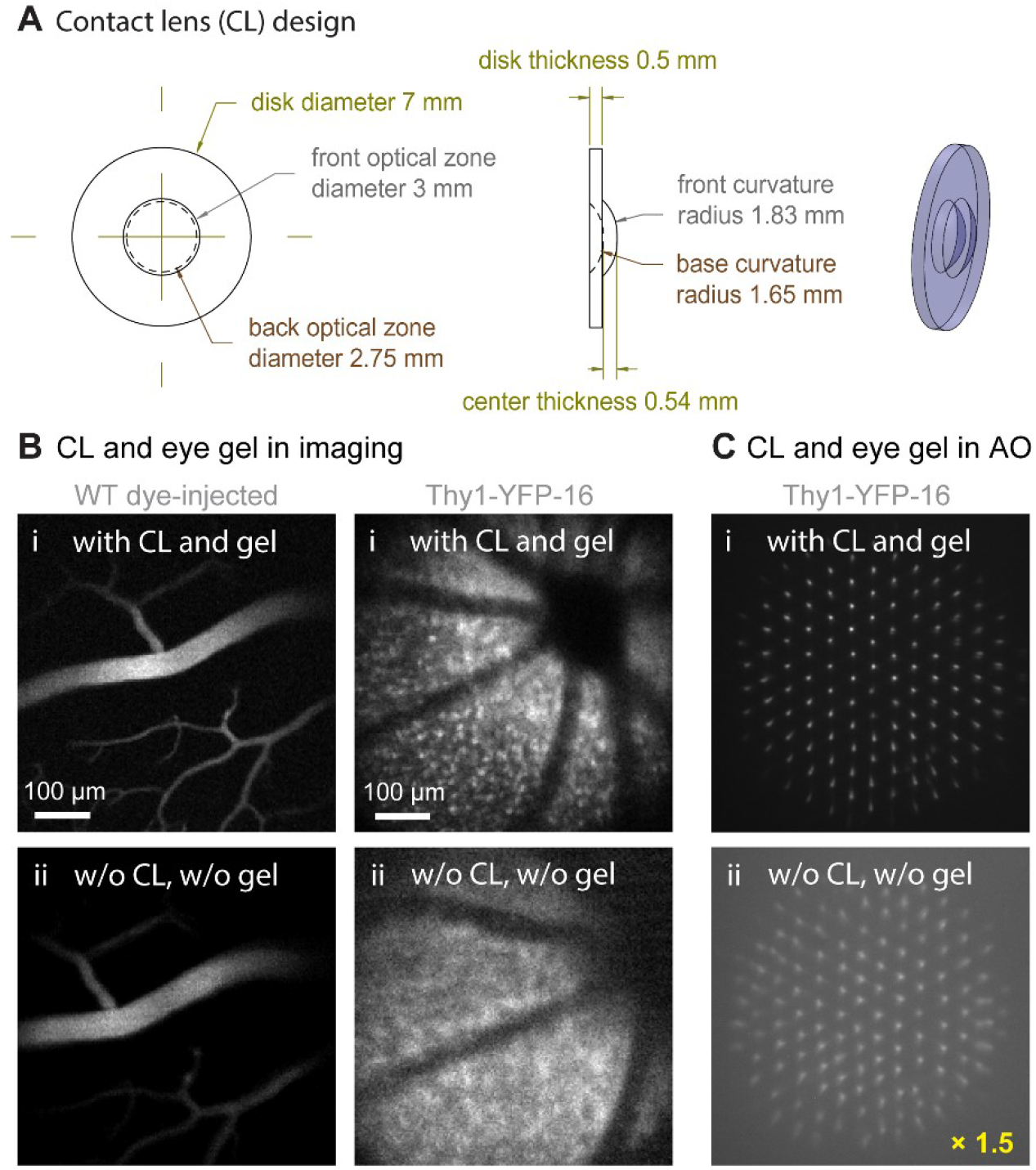
Contact lens and eye gel application improve image and wavefront sensing quality. **(A)** Design of the customized contact lens (CL). **(B)** 2PFM single-plane images of (left) retinal vasculature and (right) retinal cells acquired (i) with CL and eye gel and (ii) without CL or eye gel. All images taken with system aberration correction and normalized to (i). WT: wildtype. **(C)** Shack-Hartmann (SH) sensor images acquired from the Thy1-YFP-16 mouse retina in (B), normalized to SH image in (i). Brightness of SH image in (ii) artificially increased by 1.5× for better visualization.

**Figure S3.**
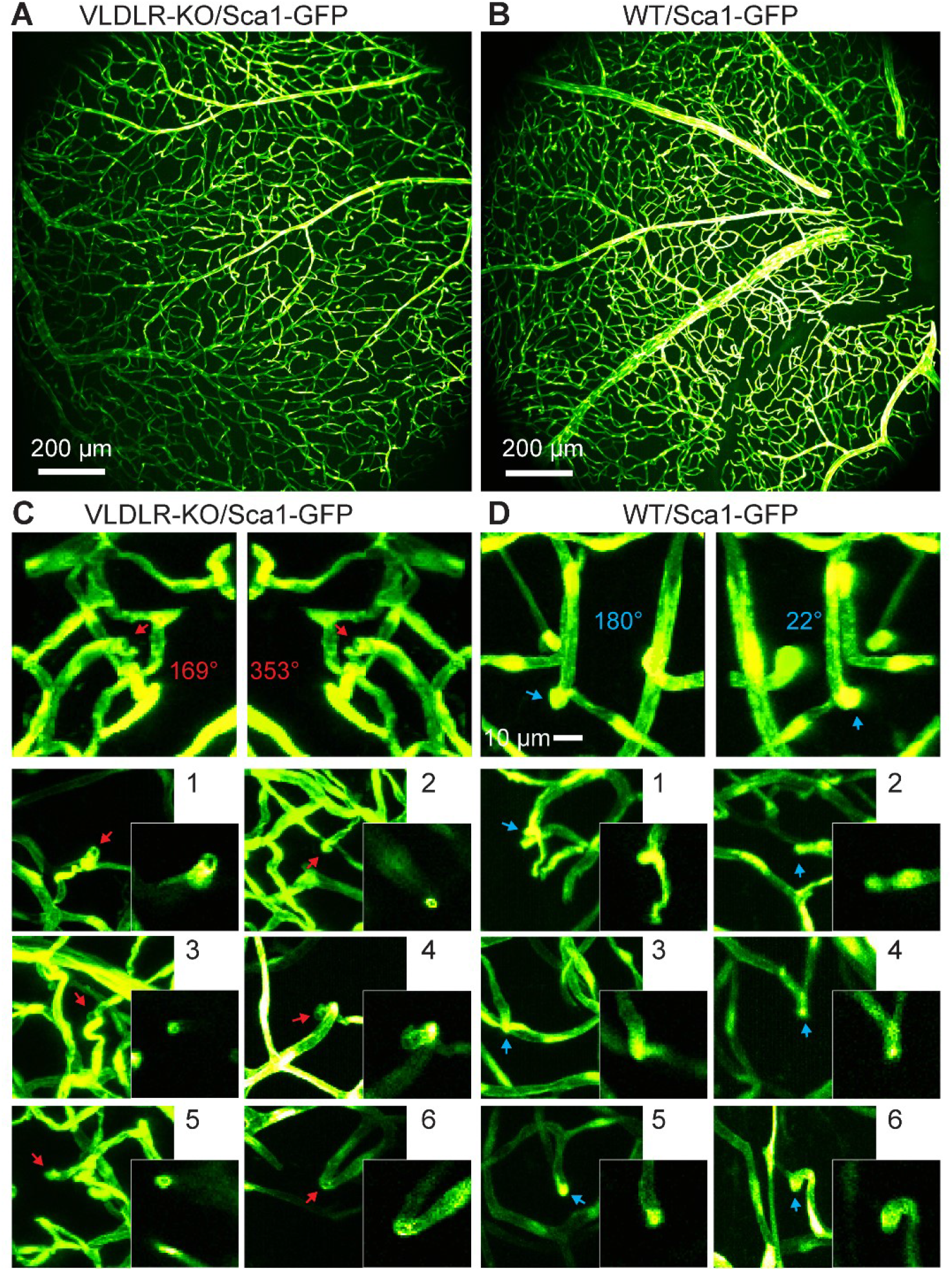
*Ex vivo* 2PFM imaging of dissected VLDLR-KO/Sca1-GFP and WT/Sca1-GFP mouse retinas. **(A**,**B)** *Ex vivo* MIPs of image stacks from **(A)** VLDLR-KO/Sca1-GFP (1380 × 1380 × 82 µm3) and **(B)** WT/Sca1-GFP (1380 × 1380 × 71 µm3) mouse retinas. **(C)** Top: 3D projected view of an example capillary lesion (red arrow) displayed at viewing angles of 169° and 353°, respectively. Bottom: MIPs of 6 more FOVs showing capillary disruptions (red arrows). Insets: single-plane zoomed-in images. **(D)** Top: 3D projected view of an example normal capillary (blue arrow) displayed at viewing angles of 180° and 22°, respectively. Bottom: MIPs of 6 more FOVs showing normal capillary structures (blue arrows). Insets: single-plane zoomed-in images. Image contrast was adjusted individually for better visualization.

**Figure S4.**
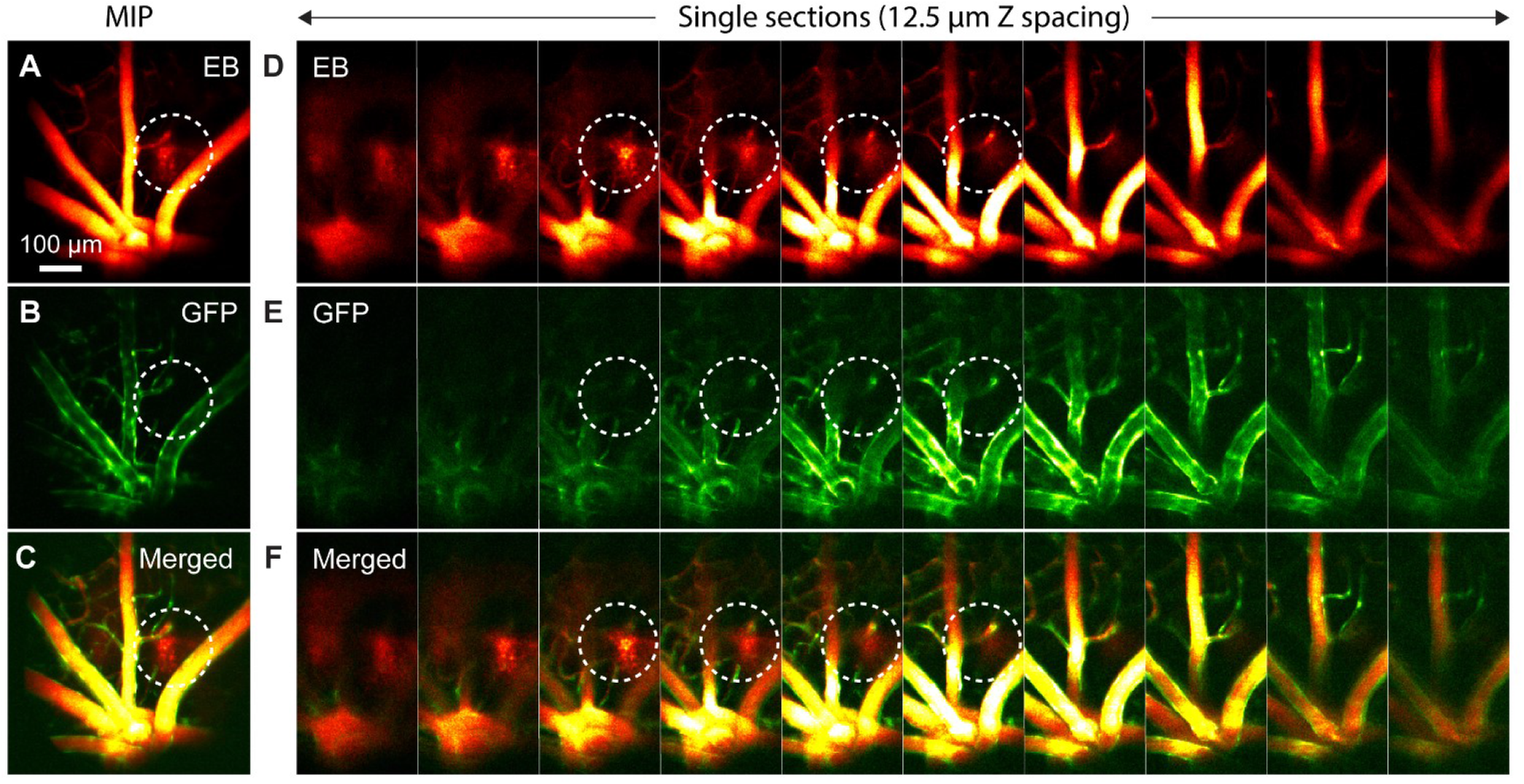
*In vivo* AO-2PFM imaging of Evans Blue (EB) leakage in healthy retina. **(A-C)** MIPs of image stacks of (580 × 580 × 130 µm3) WT/Sca1-GFP retina measured in the (**A**) near-infrared EB and (**B**) green GFP channels, and (**C**) merged images. (**D-F**) Single-plane images from (**A-C**) with 12.5 µm Z step. White dashed circles: EB leakage areas.

**Figure S5.**
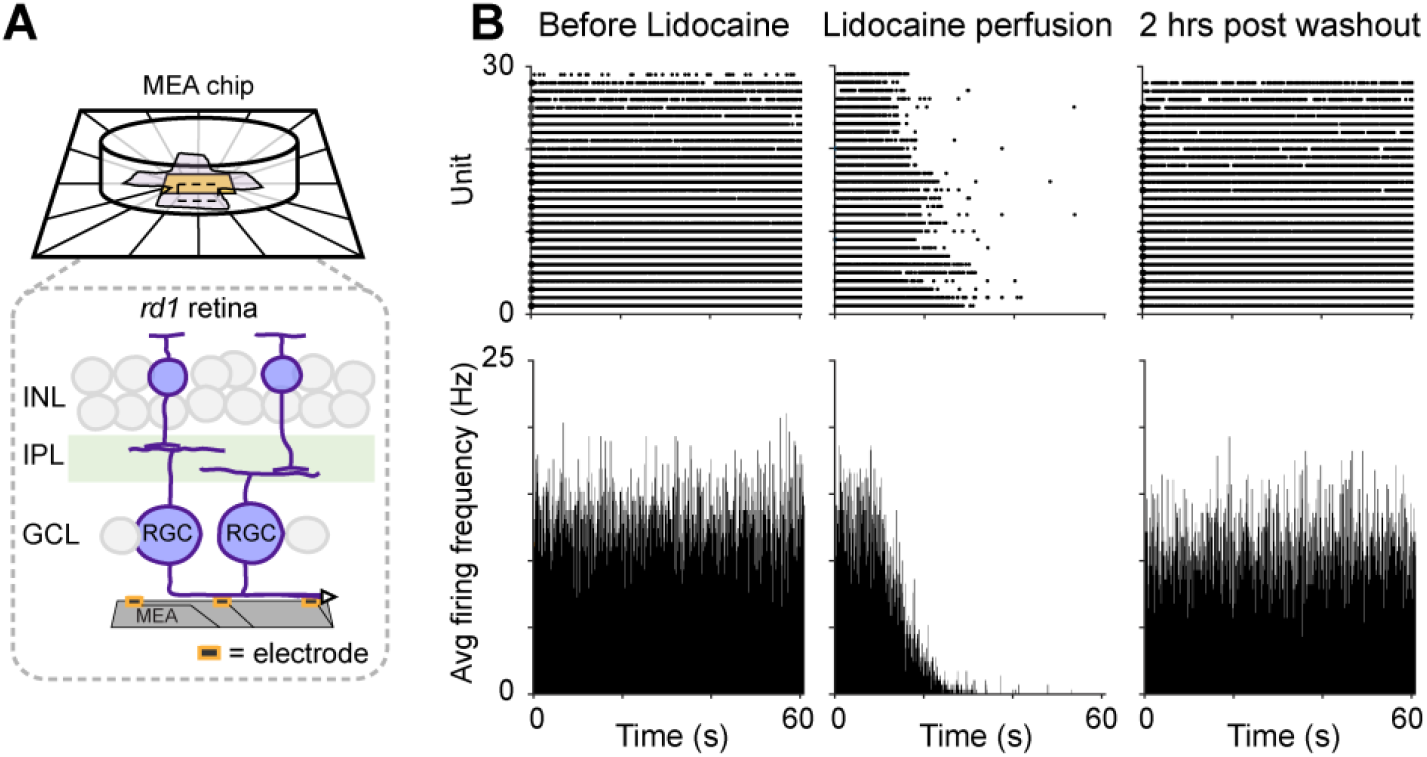
*Ex vivo* multielectrode array (MEA) recordings of Lidocaine-modified RGC hyperactivity in *rd1-*Thy1-GCaMP6s mouse retina. **(A)** MEA setup for RGC spontaneous spike activity recording. Inset: illustration of retina placement relative to MEA. **(B)** Raster and average firing frequency plots of RGCs in dissected *rd1* mouse retina (left) before and (middle) right after Lidocaine bath perfusion, and (right) 2 hours post washout, respectively.

**Figure S6.**
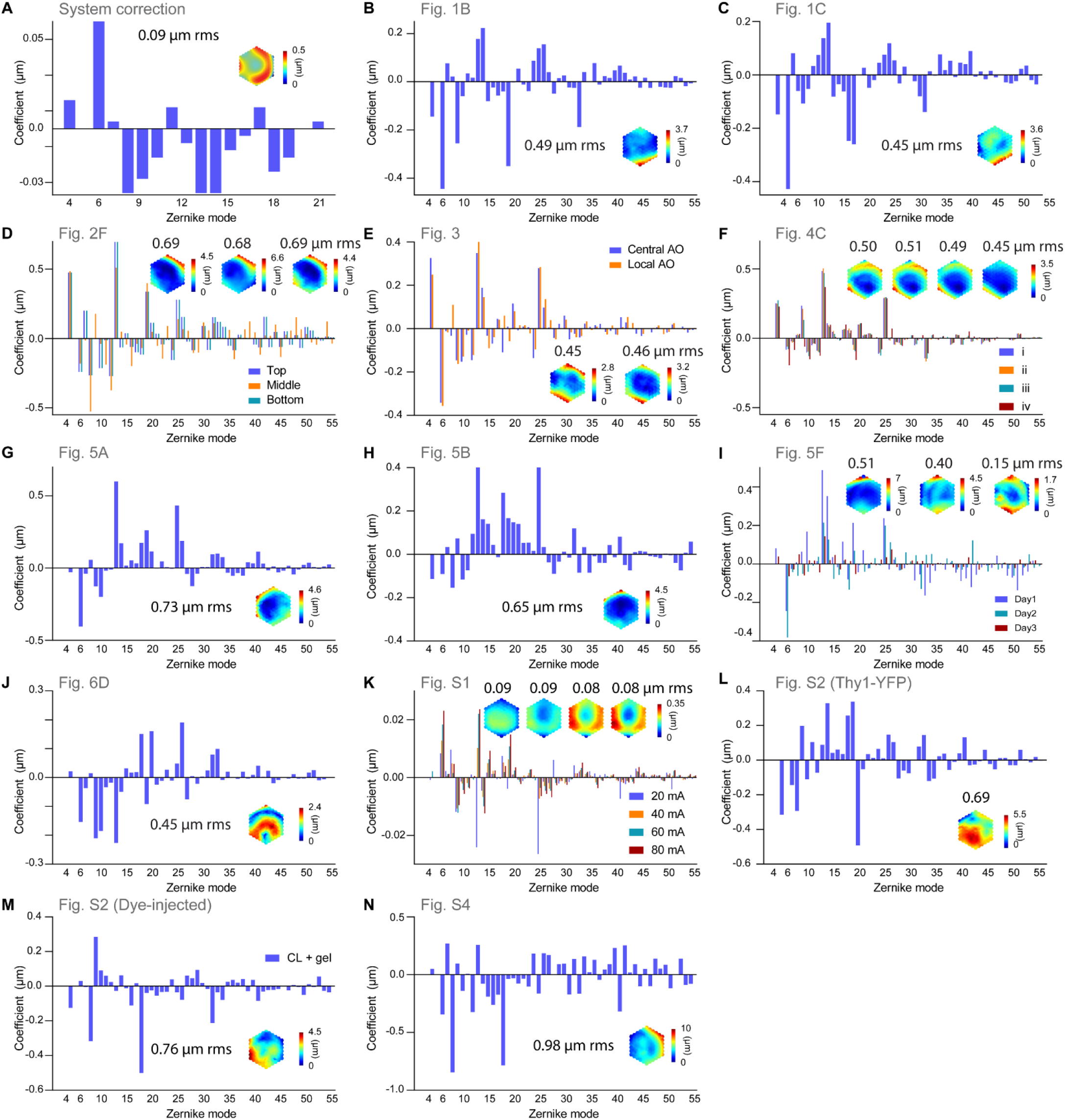
Zernike decompositions and corrective wavefronts for all experiments. All corrective wavefronts and Zernike decompositions were calculated excluding piston, tip, tilt, and defocus.

**Table S1.**
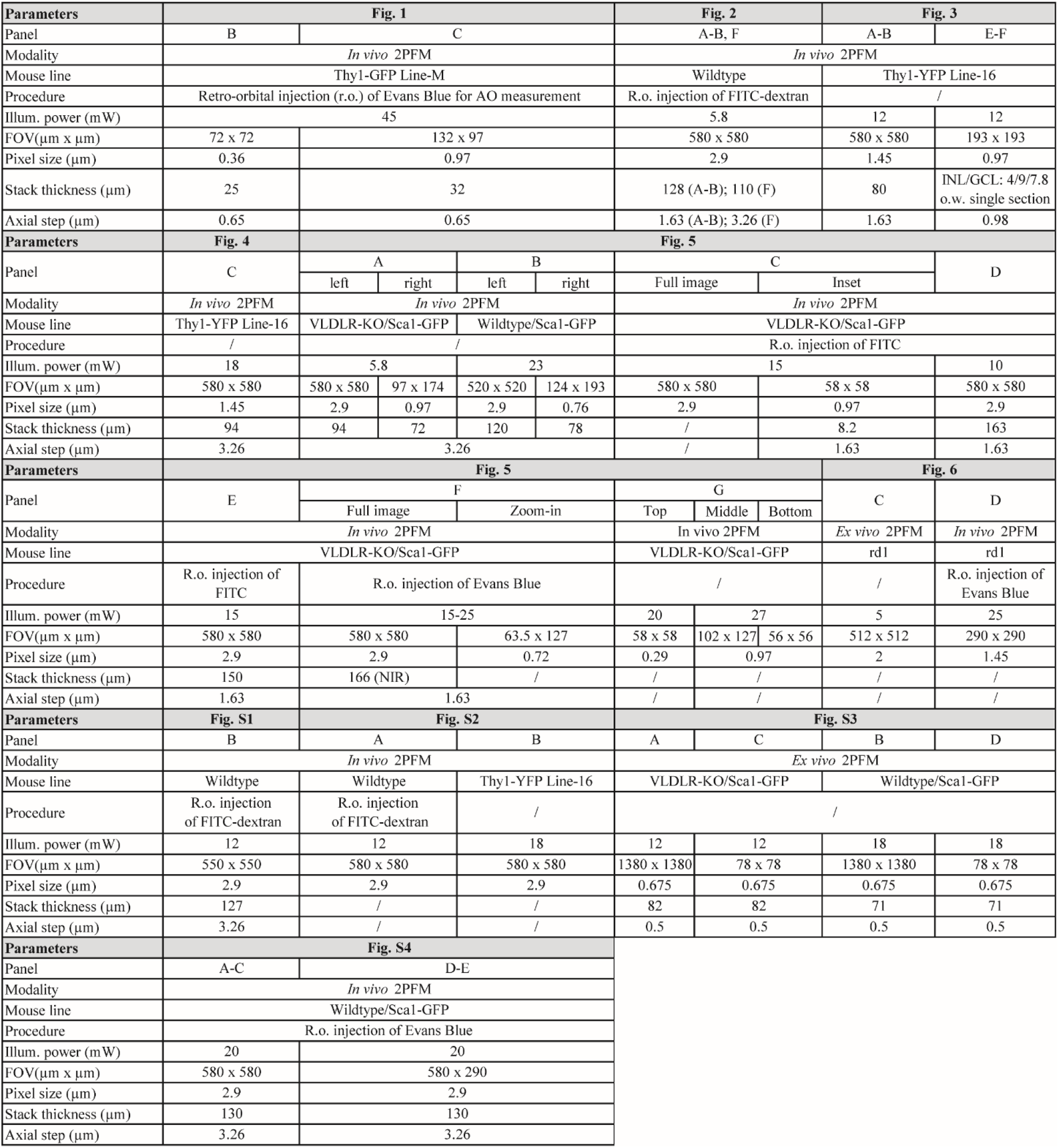
Experiment settings for all experiments.

## Notes

### Competing Interest Statement

The authors have declared no competing interest.

## References

1. Kandel, E. R., Koester, J. D., Mack, S. H. & Siegelbaum, S. A. Principles of neural science, sixth edition. (McGraw-Hill, 2021).

2. Selvam, S., Kumar, T. & Fruttiger, M. Retinal vasculature development in health and disease. Prog. Retin. Eye Res. 63, 1–19 (2018).

3. A. P. Schachat, C. P. Wilkinson, D. R. Hinton, S. R. Sadda, P. W. Ryan’s Retina. (Elsevier Health Sciences, 2017).

4. Cheung, C. Y. lui, Ikram, M. K., Chen, C. & Wong, T. Y. Imaging retina to study dementia and stroke. Prog. Retin. Eye Res. 57, 89–107 (2017).

5. London, A., Benhar, I. & Schwartz, M. The retina as a window to the brain—from eye research to CNS disorders. Nat. Rev. Neurol. 9, 44–53 (2013).

6. Lechner, J., O’Leary, O. E. & Stitt, A. W. The pathology associated with diabetic retinopathy. Vision Res. 139, 7–14 (2017).

7. Ivanova, E., Corona, C., Eleftheriou, C. G., Bianchimano, P. & Sagdullaev, B. T. Retina-specific targeting of pericytes reveals structural diversity and enables control of capillary blood flow. J. Comp. Neurol. 529, 1121–1134 (2021).

8. Jo, A. et al. Intersectional Strategies for Targeting Amacrine and Ganglion Cell Types in the Mouse Retina. Front. Neural Circuits 12, 1–15 (2018).

9. Martersteck, E. M. et al. Diverse Central Projection Patterns of Retinal Ganglion Cells. Cell Rep. 18, 2058–2072 (2017).

10. Eme-Scolan, E. & Dando, S. J. Tools and Approaches for Studying Microglia In vivo. Front. Immunol. 11, 1–10 (2020).

11. Denk, W., Strickler, J. H. & Webb, W. W. Two-Photon Laser Scanning Fluorescence Microscopy. Science 248, 73–76 (1990).

12. Euler, T., Detwiler, P. B. & Denk, W. Directionally selective calcium signals in dendrites of starburst amacrine cells. Nature 418, 845–852 (2002).

13. Baden, T. et al. The functional diversity of retinal ganglion cells in the mouse. Nature 529, 345–350 (2016).

14. Hampson, K. M. et al. Adaptive optics for high-resolution imaging. Nat. Rev. Methods Prim. 1, 68 (2021).

15. Ji, N. Adaptive optical fluorescence microscopy. Nat. Methods 14, 374–380 (2017).

16. Rodríguez, C. & Ji, N. Adaptive optical microscopy for neurobiology. Curr. Opin. Neurobiol. 50, 83–91 (2018).

17. Liang, J., Williams, D. R. & Miller, D. T. Supernormal vision and high-resolution retinal imaging through adaptive optics. J. Opt. Soc. Am. A 14, 2884 (1997).

18. Akyol, E., Hagag, A. M., Sivaprasad, S. & Lotery, A. J. Adaptive optics: principles and applications in ophthalmology. Eye 35, 244–264 (2021).

19. Palczewska, G. et al. Noninvasive two-photon microscopy imaging of mouse retina and retinal pigment epithelium through the pupil of the eye. Nat. Med. 20, 785–789 (2014).

20. Biss, D. P. et al. In vivo fluorescent imaging of the mouse retina using adaptive optics. Opt. Lett. 32, 659 (2007).

21. Alt, C., Biss, D. P., Tajouri, N., Jakobs, T. C. & Lin, C. P. An adaptive-optics scanning laser ophthalmoscope for imaging murine retinal microstructure. in Proc.SPIE vol. 7550 755019 (2010).

22. Geng, Y. et al. Adaptive optics retinal imaging in the living mouse eye. Biomed. Opt. Express 3, 715 (2012).

23. Sharma, R. et al. In vivo two-photon imaging of the mouse retina. Biomed. Opt. Express 4, 1285 (2013).

24. Wahl, D. J., Jian, Y., Bonora, S., Zawadzki, R. J. & Sarunic, M. V. Wavefront sensorless adaptive optics fluorescence biomicroscope for in vivo retinal imaging in mice. Biomed. Opt. Express 7, 1 (2016).

25. Wahl, D. J. et al. Adaptive optics in the mouse eye: wavefront sensing based vs image-guided aberration correction. Biomed. Opt. Express 10, 4757 (2019).

26. Qin, Z. et al. Adaptive optics two-photon microscopy enables near-diffraction-limited and functional retinal imaging in vivo. Light Sci. Appl. 9, 79 (2020).

27. Li, Z. et al. Fast widefield imaging of neuronal structure and function with optical sectioning in vivo. Sci. Adv. 6, eaaz3870 (2020).

28. Grulkowski, I. et al. Swept source optical coherence tomography and tunable lens technology for comprehensive imaging and biometry of the whole eye. Optica 5, 52 (2018).

29. Jian, Y., Zawadzki, R. J. & Sarunic, M. V. Adaptive optics optical coherence tomography for in vivo mouse retinal imaging. J. Biomed. Opt. 18, 056007 (2013).

30. McNabb, R. P. et al. Wide-field whole eye OCT system with demonstration of quantitative retinal curvature estimation. Biomed. Opt. Express 10, 338 (2019).

31. Wang, K. et al. Rapid adaptive optical recovery of optimal resolution over large volumes. Nat. Methods 11, 625–628 (2014).

32. Wang, K. et al. Direct wavefront sensing for high-resolution in vivo imaging in scattering tissue. Nat. Commun. 6, 7276 (2015).

33. Panagopoulou, S. I. & Neal, D. R. Zonal matrix iterative method for wavefront reconstruction from gradient measurements. J Refract Surg 21, S563–S569 (2005).

34. Feng, G. et al. Imaging Neuronal Subsets in Transgenic Mice Expressing Multiple Spectral Variants of GFP. Neuron 28, 41–51 (2000).

35. Geng, Y. et al. Optical properties of the mouse eye. Biomed. Opt. Express 2, 717 (2011).

36. Patton, N. et al. Retinal vascular image analysis as a potential screening tool for cerebrovascular disease: A rationale based on homology between cerebral and retinal microvasculatures. J. Anat. 206, 319–348 (2005).

37. Frost, S. et al. Retinal vascular biomarkers for early detection and monitoring of Alzheimer’s disease. Transl. Psychiatry 3, (2013).

38. Ikram, M. K. et al. Retinal vascular caliber as a biomarker for diabetes microvascular complications. Diabetes Care 36, 750–759 (2013).

39. Liew, G., Wang, J. J., Mitchell, P. & Wong, T. Y. Retinal vascular imaging: a new tool in microvascular disease research. Circ. Cardiovasc. Imaging 1, 156–161 (2008).

40. Bar-Noam, A. S., Farah, N. & Shoham, S. Correction-free remotely scanned two-photon in vivo mouse retinal imaging. Light Sci. Appl. 5, (2016).

41. Wang, Z., McCracken, S. & Williams, P. R. Transpupillary two-photon in vivo imaging of the mouse retina. J. Vis. Exp. 2021, 1–23 (2021).

42. Campbell, M. C. W. & Hughes, A. An analytic, gradient index schematic lens and eye for the rat which predicts aberrations for finite pupils. Vision Res. 21, 1129–1148 (1981).

43. Remtulla, S. & Hallett, P. E. A schematic eye for the mouse, and comparisons with the rat. Vision Res. 25, 21–31 (1985).

44. Wang, C. & Ji, N. Pupil-segmentation-based adaptive optical correction of a high-numerical-aperture gradient refractive index lens for two-photon fluorescence endoscopy. Opt. Lett. 37, 2001–2003 (2012).

45. Wang, C. & Ji, N. Characterization and improvement of three-dimensional imaging performance of GRIN-lens-based two-photon fluorescence endomicroscopes with adaptive optics. Opt. Express 21, 27142–27154 (2013).

46. Hoon, M., Okawa, H., Della Santina, L. & Wong, R. O. L. Functional architecture of the retina: Development and disease. Prog. Retin. Eye Res. 42, 44–84 (2014).

47. Ji, N., Sato, T. R. & Betzig, E. Characterization and adaptive optical correction of aberrations during in vivo imaging in the mouse cortex. Proc. Natl. Acad. Sci. 109, 22–27 (2012).

48. Wang, C. et al. Multiplexed aberration measurement for deep tissue imaging in vivo. Nat. Methods 11, 1037–1040 (2014).

49. Sun, W., Tan, Z., Mensh, B. D. & Ji, N. Thalamus provides layer 4 of primary visual cortex with orientation- and direction-tuned inputs. Nat. Neurosci. 19, 308–315 (2016).

50. Yannuzzi, L. A. et al. Retinal angiomatous proliferation in age-related macular degeneration. Retina 21, 416–34 (2001).

51. Heckenlively, J. R. et al. Mouse model of subretinal neovascularization with choroidal anastomosis. Retina 23, 518–522 (2003).

52. Hu, W. et al. Expression of VLDLR in the retina and evolution of subretinal neovascularization in the knockout mouse model’s retinal angiomatous proliferation. Investig. Opthalmology Vis. Sci. 49, 407 (2008).

53. Li, C. et al. Biochemical Alterations in the Retinas of Very Low-Density Lipoprotein Receptor Knockout Mice. Arch Ophthalmol. 125, (2007).

54. Xia, C., Lu, E., Zeng, J. & Gong, X. Deletion of LRP5 in VLDLR knockout mice inhibits retinal neovascularization. PLoS One 8, e75186 (2013).

55. Chen, T.-W. et al. Ultrasensitive fluorescent proteins for imaging neuronal activity. Nature 499, 295–300 (2013).

56. Chang, B. et al. Retinal degeneration mutants in the mouse. Vision Res. 42, 517–525 (2002).

57. Sekirnjak, C. et al. Changes in physiological properties of rat ganglion cells during retinal degeneration. J. Neurophysiol. 105, 2560–2571 (2011).

58. Telias, M. et al. Retinoic Acid Induces Hyperactivity, and Blocking Its Receptor Unmasks Light Responses and Augments Vision in Retinal Degeneration. Neuron 102, 574-586.e5 (2019).

59. Telias, M. et al. Retinoic acid inhibitors mitigate vision loss in a mouse model of retinal degeneration. Sci. Adv. 8, 1–18 (2022).

60. Cao, K. J. et al. Cyclodextrin-Assisted Delivery of Azobenzene Photoswitches for Uniform and Long-Term Restoration of Light Responses in Degenerated Retinas of Blind Mice. Adv. Ther. 4, (2021).

61. Liu, X. et al. Effect of contact lens on optical coherence tomography imaging of rodent retina. Curr. Eye Res. 38, 1235–1240 (2013).

62. Ikeda, W., Nakatani, T. & Uemura, A. Cataract-preventing contact lens for in vivo imaging of mouse retina. Biotechniques 65, 101–104 (2018).

63. Zhang, Q., Pan, D. & Ji, N. High-resolution in vivo optical-sectioning widefield microendoscopy. Optica 7, 1287 (2020).

64. Jia, Y. et al. Quantitative optical coherence tomography angiography of vascular abnormalities in the living human eye. Proc. Natl. Acad. Sci. 112, E2395–E2402 (2015).

65. Hormel, T. T. et al. Plexus-specific retinal vascular anatomy and pathologies as seen by projection-resolved optical coherence tomographic angiography. Prog. Retin. Eye Res. 80, 100878 (2021).

66. Prahst, C. et al. Mouse retinal cell behaviour in space and time using light sheet fluorescence microscopy. Elife 9, 1–29 (2020).

67. Sulai, Y. N. & Dubra, A. Non-common path aberration correction in an adaptive optics scanning ophthalmoscope. Biomed. Opt. Express 5, 3059 (2014).

68. Booth, M. J. Adaptive optical microscopy: the ongoing quest for a perfect image. Light Sci. Appl. 3, e165 (2014).

69. Schindelin, J. et al. Fiji: an open-source platform for biological-image analysis. 9, 676–682 (2012).

